# Ultrastructural membrane dynamics of mouse and human cortical synapses

**DOI:** 10.1101/2024.12.26.630393

**Authors:** Chelsy R. Eddings, Minghua Fan, Yuuta Imoto, Kie Itoh, Xiomara McDonald, Jens Eilers, William S. Anderson, Paul F. Worley, Kristina Lippmann, David W. Nauen, Shigeki Watanabe

## Abstract

Live human brain tissues provide unique opportunities for understanding synaptic transmission. Investigations have been limited to anatomy, electrophysiology, and protein localization—while crucial parameters such as synaptic vesicle dynamics were not visualized. Here we utilize zap-and-freeze time-resolved electron microscopy to overcome this hurdle. First, we validate the approach with acute mouse brain slices to demonstrate that slices can be stimulated to produce calcium signaling. Next, we show that synaptic vesicle endocytosis is induced in both mouse and human brain slices. Crucially, clathrin-free endocytic pits appear immediately next to the active zone, where ultrafast endocytosis normally occurs, and can be trapped at this location by a dynamin inhibitor. In both species a protein essential for ultrafast endocytosis, Dynamin 1xA, localizes to the region peripheral to the active zone, the putative endocytic zone, indicating a possible conserved mechanism between mouse and human. This approach has the potential to reveal dynamic, high-resolution information about synaptic membrane trafficking in intact human brain slices.

## Introduction

Direct investigation of synaptic transmission in human brain tissues will increase our understanding of typical brain states and how they are impacted by age and disease. Dynamic measures of membrane properties and firing patterns of human neurons are observed via electrophysiology of *ex vivo* brain slices. Electrophysiology enables quantitative evaluation of synaptic properties of specific neuronal subtypes in particular brain regions. For example, human neurons located in different cortical layers exhibit heterogeneous morphologies, passive membrane and suprathreshold properties, as well as firing rates—with layer 5 pyramidal neurons presenting higher frequency firing rates compared to layer 2/3 and layer 3c^1^. These properties can also change across different time scales. For instance, the resting membrane potential and resistance of layer 2/3 pyramidal neurons differ from infancy to old age^2^. Consequently, electrophysiology provides information on quantal content, evoked and spontaneous vesicular release, and the plastic nature of synaptic transmission. Likewise, this method reveals species-specific differences in synaptic reliability and circuit connectivity between mice and humans. Human cortical pyramidal synapses are more reliable and less likely to fail compared to mouse pyramidal synapses—with a failure rate of 0% compared to 25% in mice^3^. Work on human CA3 hippocampal tissues suggest higher temporal precision in human synapses compared to rodent synapses. Human hippocampal synapses show pronounced adaptation, more narrow interspike intervals and overall lower synaptic fluctuations^4^. Yet, relying solely on electrophysiology cannot easily discern the anatomical and structural features of synapses.

Cellular and synaptic morphologies are often characterized by electron microscopy (EM). Enhanced resolution clearly distinguishes presynaptic from postsynaptic compartments and reveals the relative locations of organelles. An advantage of EM is that it yields non-targeted information about the morphology of all cells in a tissue while preserving their spatial relationships^5^. Moreover, electron micrographs provide detailed information about how neurological diseases impact non-neuronal cells and myelination in patient brains. For example, post-mortem Alzheimer’s patient hippocampal tissues contain ‘dark astrocytes’ showing signs of cellular stress like altered mitochondria and electron-dense plaques^6^. On the other hand, examinations of post-mortem Multiple Sclerosis patient brains show reduced myelination and possible compensatory axonal swelling in regions of normal-appearing white matter^7^. Along with the advent of volumetric imaging and machine-learning based segmentation approaches, millions of human synaptic connections are now being mapped. Recently a petascale dataset was published from a 1 mm^3^ human cortical biopsy illustrating vasculature, rare multisynaptic connections, and intact surrounding tissue^8^. Other EM datasets reveal subtle connections between human neurons and glia through cilia-based contacts^9^, and a 10-fold expansion of the interneuron-to-interneuron network in humans compared to mice^10^. Overall, ultrastructural investigations of human brain tissues provide additional insights into how synaptic circuits are situated within the larger cerebral context. However, electron micrographs are static images of tissue and do not provide information on synaptic release properties beyond the distribution of organelles.

These various methods of studying synapses can require vastly different sample preparation requirements, resulting in a frequent ‘structure-function’ gap in understanding synaptic transmission. Time-resolved electron microscopy is situated as a potential resolution to this structure-function dilemma. Combining optogenetic stimulation and high-pressure freezing ‘flash-and-freeze’ EM has been successfully used in: *C. elegans* neuromuscular junctions^11^, cultured mouse hippocampal neurons^12^, acute mouse brain slices^13^, and mouse hippocampal organotypic slice cultures^14^. An advantage and restriction of flash-and-freeze is its reliance on channelrhodopsin activation of neurons. Channelrhodopsin precisely activates select circuits through restricted expression to a subset of neurons^13,14^. This exogenous activation system can limit users to transgenic mouse lines or virally infected tissues. In addition, the number of elicited action potentials in each neuron may be variable due to differential expression levels^12^.

Although human organotypic slice cultures can be transduced with channelrhodopsin expression^15^, quality long-term culture likely requires readily accessible human cerebrospinal fluid sources^16^, which can be difficult to obtain. Increased culture times can result in glial scaring at the tissue surface as well as slight tissue flattening (∼20% for human brain)^16^ and decreased tissue stability, which must be accounted for in subsequent experiments. On the contrary, acute brain slices offer near *in vivo* tissue states for rapid analysis of synaptic properties^17^. As such acute slices offer a platform for more samples to be tested without the need for extended culturing times. Being able to investigate a higher number of brain tissues from different individuals offers a more comprehensive understanding of the human brain—expanding the extent of results from a single patient to multiple people and possibly revealing conserved cellular processes. To push methods towards obtaining ultrastructural information from activated synapses across several humans with more native cytoarchitectures, a rapid approach suitable for acutely resected brain tissue is necessary.

Here, we adapt ‘zap-and-freeze’ EM for use with acute brain slices as another modality to bridge form and function, yielding functional and ultrastructural information about presynaptic membrane trafficking. Zap-and-freeze EM uses transient electric field stimulation to activate neurons before high-pressure freezing^18^. By controlling the stimulation paradigm as well as the time interval between stimulation and freeze, snapshots of dynamic events are captured with millisecond temporal resolution. We applied zap-and-freeze EM to acute mouse brain slices and acute human neocortical slices obtained from epilepsy surgeries. We demonstrate that synaptic activities can be induced in slices loaded on the zap board for a high-pressure freezer. Our results indicate that synaptic vesicle recycling in human neocortical presynapses can be mediated by ultrafast endocytosis, owing to a depot of the protein Dynamin 1xA (Dyn1xA), similar to mouse hippocampal synapses. Our approach can enable ultrastructural characterizations of activity-dependent membrane trafficking at synapses in acute brain slices.

## Results

### The ‘zap board’ induces calcium signaling in acute mouse brain slices

Towards our goal of visualizing synaptic vesicle dynamics in human tissues, we began by determining if the microcircuit chip used in zap-and-freeze (called the ‘zap board’ hereafter) is compatible with brain slices. The board’s circuit diagram and detailed electrical properties are described in Figure S1A-I. Originally, the zap board was constructed to apply a 10 V/cm electric field to induce action potential-driven synaptic activities in cultured mouse neurons^19^. Provided the orientation of axons is highly variable in culture, most neurons within the electric field can be stimulated^19^. Considering an electric field is a vector with both magnitude and direction we were unsure if tissue directionality within the zap board was important, since the orientation of axons in an intact neural circuit may influence the generation of action potentials in slices^20,21^. Therefore, we first tested the zap board using axons with a well-defined circuit orientation in slices: the parallel fibers of granule cells, which form synapses with Purkinje cells, in the mouse cerebellum (Figure 1A).

**Figure 1.**
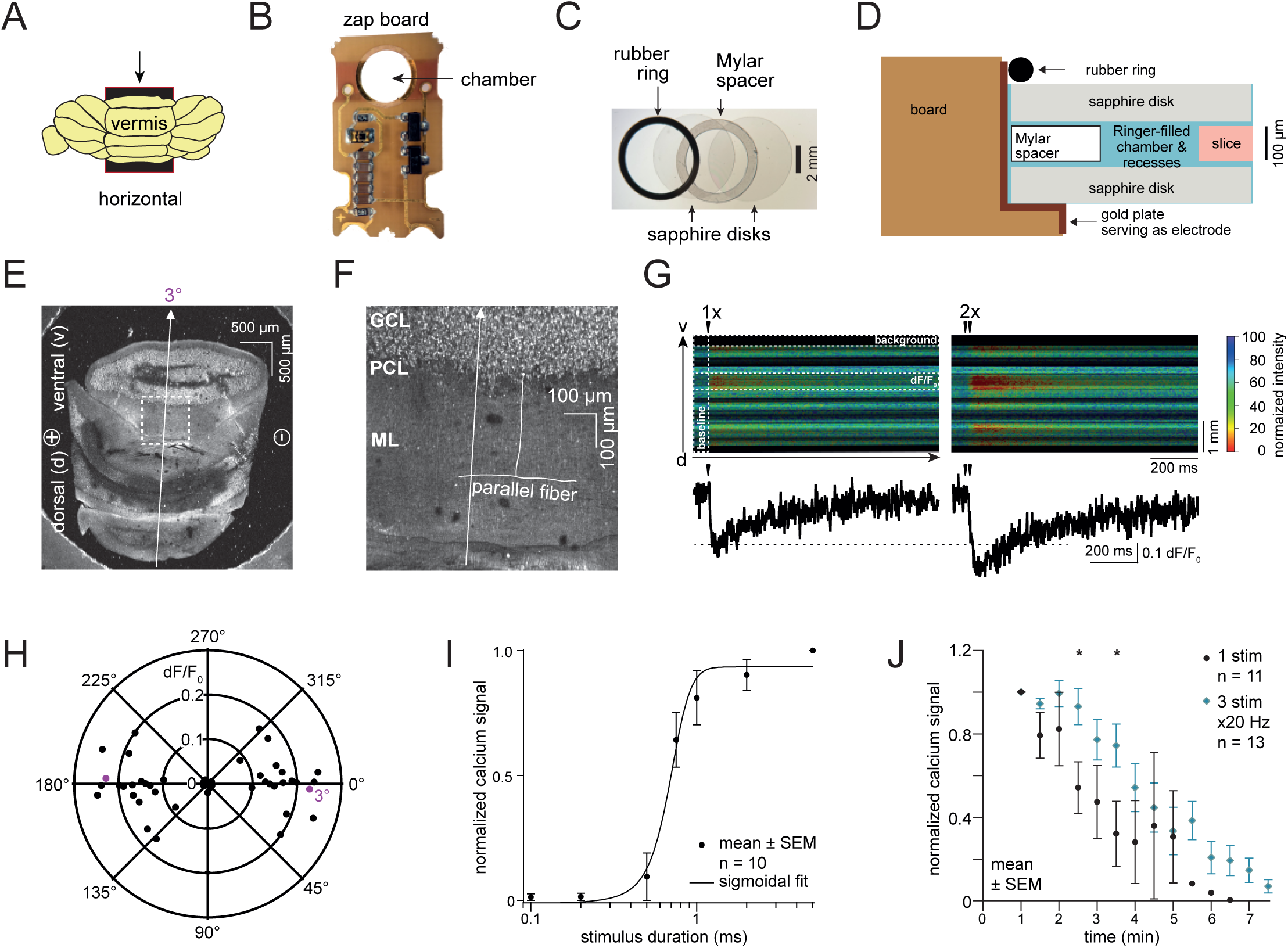
2-photon calcium imaging of acute mouse cerebellar slices on the zap board. (A) Acute mouse cerebellar slices were prepared by sectioning the cerebellar vermis horizontally to produce axons with known orientation. Arrow indicates vibratome cut direction. (B) A zap board enabling electrical stimulation in the Leica EM ICE high-pressure freezer. (C) Overlay image of a 6 mm wide rubber O-ring, two sapphire disks and a Mylar spacer ring in between two sapphire disks. When assembled, a specimen is sandwiched between the two sapphire disks. (D) A schematic showing another view of the assembly holding the slice in the zap board chamber. (E) Horizontal 100 µm cerebellar slice lying between two sapphire discs and a Mylar spacer ring for 2-photon imaging on the zap board. Parallel fibers (PF) from the ascending axons of granule cells (GCs) are in line with the electrical dipoles ((-) to (+)) and were imaged orthogonally in the line-scan mode (arrow, 3° deviated in this example). (F) Zoom-in image of a slice indicating the PFs in the molecular layer (ML), arising from GCs in the granule cell layer (GCL). An arrow depicts the orientation of the line scan. PCL: Purkinje cell layer. (G) Normalized Fura-2, AM signal after a single (1x) and two consecutive (2x, 50 ms apart) stimuli. (H) Polar coordinate plot of normalized Ca^2+^ signal vs. linescan / PF orientation. 0/180° account for PFs lying in line with the electrical dipole, while at 90/270° PFs lie orthogonally to the dipole. Circles indicate Ca^2+^ signal strength. n=25 slices from 4 mice. (I) Plot showing the normalized Ca^2+^ signal versus stimulus duration. Data are presented as mean ± SEM. n=10 slices from 2 mice. (J) Plot showing the normalized peak Ca^2+^ signal versus time after assembling the cerebellar slice in the zap board chamber for one stimulus and three stimuli at 20 Hz. Data are presented as mean ± SEM. n=11 slices from 2 mice (1 stimulus, black markers) and n=13 slices from 3 mice (3 stimuli, blue markers). Stars indicate a significant difference (Mann-Whitney U test, p = 0.02 for 2.5 minutes and p = 0.04 for 3.5 minutes).

To visualize neuronal activity, we performed 2-photon calcium imaging in acute mouse cerebellar slices. Slices were placed between two sapphire disks, separated by a 100 µm Mylar spacer ring within the zap board (Figure 1B-D). The zap board is normally activated by pulses of blue light of various durations^19^ (typically 1 ms is sufficient to induce an action potential in cultured neurons^22^). To avoid blue light interference during 2-photon imaging when activating the zap board, we relocated the photodiode to an external optocoupler and drove the board with transistor-transistor logic (TTL) signals from a patch-clamp amplifier. In this setup, the zap board remains optically activated with blue light, as it is in the high-pressure freezer, but can now be placed on a 2-photon laser-scanning microscope (Figure S1). After determining the optimal stimulus parameters (e.g., 2.55 V LED command voltage) in this system with external resistances (Figure S1), we tested the electrical stimulation on slices.

For calcium imaging, 100 µm mouse cerebellar slices were incubated with the membrane-permeable calcium indicator Fura-2, AM (which darkens with increasing calcium concentrations) and mounted on the zap board (Figure 1B-D). First, we oriented cerebellar slices such that the parallel fibers were themselves parallel with the electric field (3°, Figure 1E,F). After applying a single command voltage to the LED to activate the zap board for 1 ms, calcium transients were successfully detected in the molecular layer (ML; Figure 1G). Calcium signaling was enhanced when two consecutive 1 ms pulses were applied 50 ms apart (Figure 1G), indicating synaptic activity in parallel fibers^23^ is successfully induced by the zap board. To test the effect of neural circuit orientation within the zap board, we varied the angular orientation of brain slices relative to the device’s electric dipoles. Calcium imaging suggested that neuronal signaling was greatest when the parallel fibers were in line with the electric dipole (0°) and decreased progressively as the angle changed to 45° (Figure 1H). Beyond 45°, essentially no calcium signal was detected (dF/F_0_ close to 0, Figure 1H), suggesting that the parallel fibers were not sufficiently stimulated. Interestingly at 90°/270° where ascending axons should be in line with the electric field, no calcium signal was observed—likely because superficial parallel fibers were imaged, whose ascending axons were cut during the slicing procedure. Nonetheless, these data suggest brain slices can be activated on the zap board with a single 1 ms pulse.

Next, we varied the electric pulse duration and the incubation time of the slice in the chamber of the zap board, while keeping the slice at close to 0° relative to the dipoles. When an electric pulse was applied for 0.1-0.5 ms, which is typically used in electrophysiology experiments^24–26^, almost no signal was detected (Figure 1I). The calcium signal nearly peaked with a 1 ms pulse (Figure 1I), suggesting that a 1 ms electrical pulse is sufficient for action potential induction or, at least, the detection of calcium signals in this setup. Finally, to address how long slice health can be maintained on the zap board, we measured the calcium signals every 30 s over 7 min after a top sapphire disk was placed over the brain slice (Figure 1C,D, and J). For the first 2 min, calcium signaling was maintained. However, beyond 2 min signals drastically decreased, becoming undetectable after a single stimulus by 6 min (black markers, Figure 1J). We also applied multiple stimuli to assess if the observed tissue deterioration was dependent on the number of stimuli given over time. When a train of three stimuli was applied at 20 Hz, the signal also declined within 2 to 3 min (blue markers, Figure 1J). Yet, this decline after multiple stimuli was less pronounced compared to a single stimulus—with signals remaining detectable for up to 7 min, an effect presumably due to a higher signal-to-noise ratio of train-induced calcium signals. Thus, for subsequent ultrastructural experiments we applied a 1 ms electric field pulse to the specimen and discarded all specimens not frozen within 2 min after they were enclosed within two sapphire disks.

As many available live human brain tissues are neocortical, we also prepared and tested mouse cortical slices to confirm that the zap board could stimulate activity in the relevant tissue type (Figure S1J-L). Unlike the more strict axon orientation of the cerebellum, cortical regions contain dispersed axons spanning within and across layers. For example, layer 5 pyramidal neurons in the mouse cortex extend axons locally within layer 5, synapsing onto inhibitory microcircuits, and also distally synapse onto layer 6 excitatory neurons^27^. It is also suggested that axons from acutely resected human brain tissues can display directionality based on their location within the cortex—axons in deeper layers appearing more columnar with directional bias, while axons in superficial layers are more randomized^28^. As such we did not change the angular orientation of cortical slices and simply tested for activity using the previously determined parameters. To help visually maintain some consistency across samples, cortical somatic layers were laid parallel with the electric field, and a line scan was performed perpendicular across the layers (Figure S1J). With a single electric field stimulus, we successfully stimulated calcium signaling in the cortex, and calcium signals increased with an increasing number of stimuli at 20 Hz (Figure S1K,L). Therefore, these results suggest that the zap board can activate cortical brain slices.

### Visualizing synaptic vesicle endocytosis in acute mouse brain slices

As human brain samples are rare and precious, we initially explored membrane dynamics using zap-and-freeze in acute mouse cortical slices. Following slicing and a slightly modified NMDG recovery^29^ (see Methods), we mounted each brain slice such that the cortical somatic layers were parallel to the zap board’s electric field to maintain some visual consistency across samples. A single 1 ms stimulus was given, and individual slices frozen at 100 ms, 1 s and 10 s. Unstimulated slices were used as controls. The selected time points were previously used to visualize synaptic vesicle recycling in cultured hippocampal neurons^12,30–34^ and can potentially capture several endocytic pathways (including kiss-and-run, ultrafast endocytosis, and clathrin-mediated endocytosis)^12,35^. Fluid phase makers of endocytosis such as cationized ferritin^12,34^ were not used on brain slices since their tissue penetration would not be sufficiently uniform. Tissue gross morphologies were left largely intact (Figure 2A) when passed through the high-pressure freezer using 15% polyvinylpyrrolidone (PVP) as a cryoprotectant—chosen because of its past success in flash-and-freeze^13^. Similar to dissociated hippocampal neurons and slices, mouse cortical axons exhibited a pearled morphology^36^ within acute slices (Figure 2A i and ii). Synapses were also readily visible (Figure 2A iii and iv).

**Figure 2.**
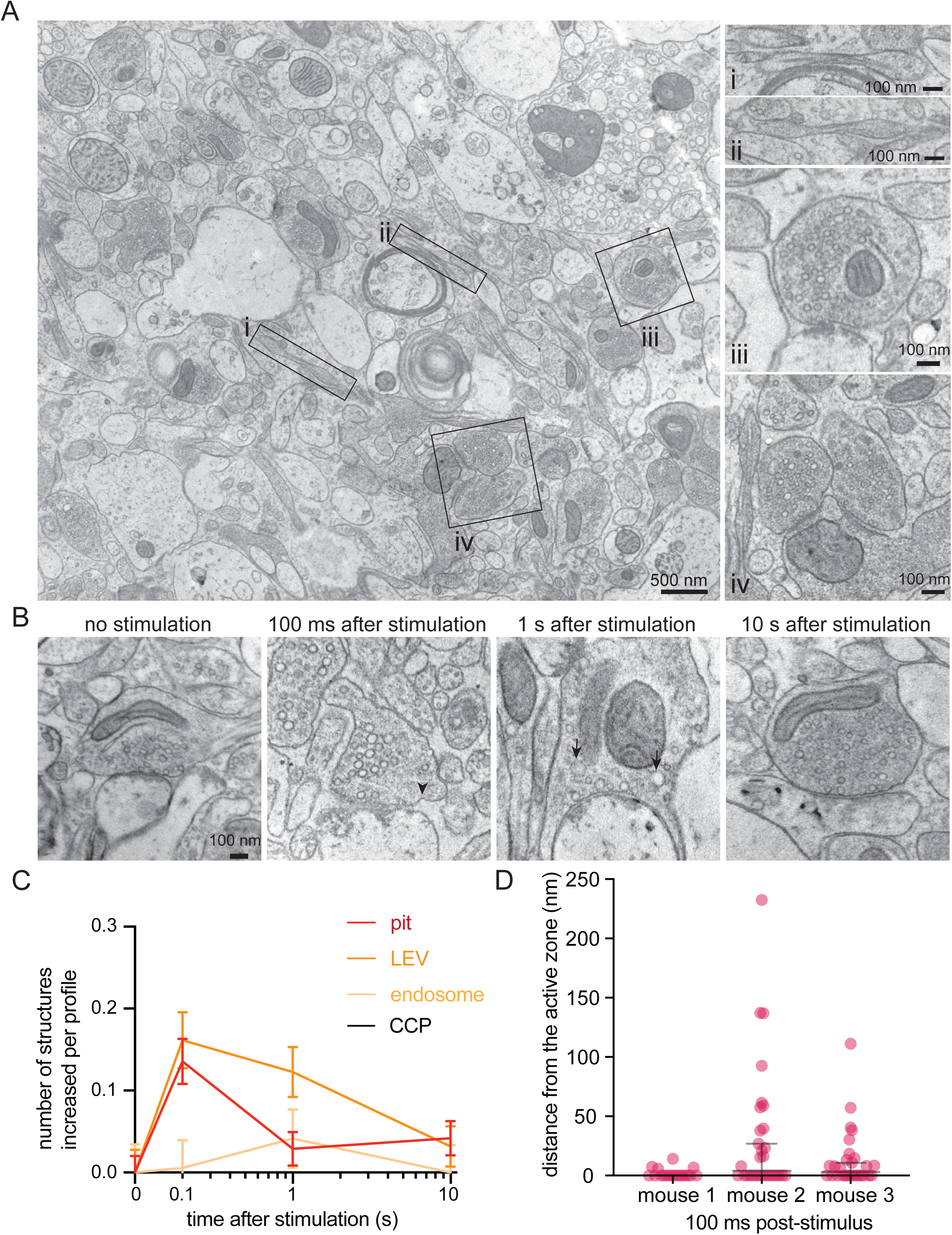
Ultrafast endocytosis in cortical synapses of acute mouse brain slices. (A) Low magnification overview of an acute mouse brain slice visualized with zap-and-freeze EM (from an unstimulated slice). Axons with pearled morphologies are highlighted as panels (i and ii), along with examples of synapses (iii and iv). (B) Example electron micrographs showing endocytic pits (black arrowheads) and putative large endocytic vesicles (LEV) and endosomes (black arrows) at the indicated time points in cortical regions of acute mouse brain slices. More example TEM images are provided in Figure S2. (C) Plots showing the increase in number of each endocytic structure per synaptic profile after a single stimulus. Data are pooled from three experiments and presented as mean ± SEM. CCP: clathrin-coated pits. See Data Table S1 for n values and detailed numbers for each time point. (D) Plot showing the distance distribution of putative uncoated endocytic pits from the edge of an active zone 100 ms post-stimulus in acute slices from n=3 mice. Data are presented as median ± 95% confidence interval. Each dot represents a pit.

Ultrastructural analysis of stimulated synapses revealed apparent induction of the synaptic vesicle cycle (Figure 2B,C). At 100 ms post-stimulus, uncoated pits appeared at putative ultrafast endocytosis sites—primarily within a range of 0-4 nm from the edge of an active zone, defined as the membrane region opposite the postsynaptic density^12,34^ (Figure 2D; median pit distances: 0, 3.892, and 3.142 nm for mouse 1, 2, and 3 respectively). These uncoated pits at 100 ms were typically shallow with diameters ranging from 35-39 nm and neck widths ranging from 58-72 nm (Figure S2A-C; median pit diameters: 38.031, 39.908, and 35.542 nm; median pit neck widths: 69.897, 72.619, and 58.523 nm for mouse 1, 2, and 3 respectively). When testing was repeated in two additional mice, uncoated pits also formed at 100 ms within 3-32 nm from an active zone (Figure S2E median pit distances: 32.354 and 3.326 nm for mouse 4 and 5 respectively), with similar diameters and neck widths (Figure S2A-C; median pit diameters: 31.816 and 35.244 nm; median pit neck widths: 58.157 and 60.219 nm for mouse 4 and 5 respectively). These pits likely represent endocytic structures rather than exocytic intermediates because they remained trapped on the plasma membrane when slices from these final two mice were treated with 80 µM Dynasore^37^, a non-specific endocytosis inhibitor^38^, for 2 min prior to zap-and-freeze testing (Fig. 3A,C). Furthermore, pre-treatment of slices with 1 µM tetrodotoxin (TTX) for 10 min nearly abolished the pits’ prevalence (Fig. 3A,B), suggesting that the formation of these structures is activity-dependent.

**Figure 3.**
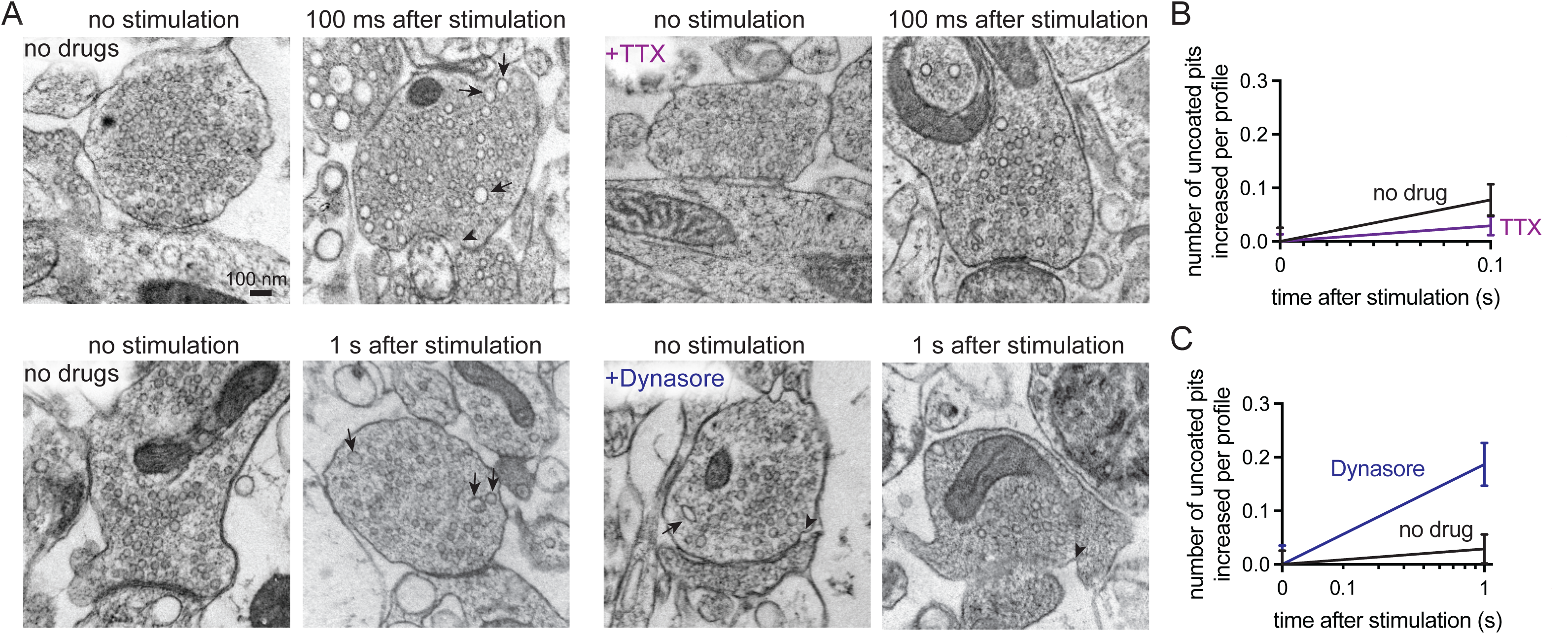
Uncoated pits in acute mouse cortical slices are activity-dependent and Dynasore-sensitive. (A) Example electron micrographs showing endocytic pits (black arrowheads) and putative large endocytic vesicles (LEV) at the indicated time points in acute brain slices from n=2 mice, given: no drugs, tetrodotoxin (TTX), or Dynasore. (B) Plot of no drug versus TTX treated slices, comparing the increase in number of uncoated pits per synaptic profile, 100 ms after a single stimulus. Data are presented as mean ± SEM. (C) Plot of no drug versus Dynasore treated slices, comparing the increase in number of uncoated pits per synaptic profile, 1 s after a single stimulus. Data are presented as mean ± SEM. See Data Table S1 for n values and detailed numbers for each time point.

Further analysis revealed that these uncoated pits are likely internalized over time. The number of large vesicles (diameter over 60 nm) also peaked at 100 ms, followed by a slight increase in the number of endosomes (spherical organelles with diameter 100 nm or more, or oblong organelles larger than synaptic vesicles) 1 s post-stimulus (Figure 2B,C; Figure S3A-C for additional micrographs). The size of endosomes ranged from 5,851-7,775 nm^2^ (Figure S2D; median endosome sizes: 6,463.22, 6,903.58, 5,850.65, 7,774.78, and 6,158.98 nm^2^ for mouse 1, 2, 3, 4, and 5 respectively). No apparent clathrin-coated pits or Ω-structures indicative of potential kiss-and-run like events^39–41^ formed at synapses at any of the tested time points (Figure 2C; see Figure S3D for micrographs of uncoated pits versus one observed clathrin-coated pit on a dendrite from this dataset for comparison). These results suggest that ultrafast endocytosis likely occurs in cortical synapses after a single stimulus.

To verify that the observed uncoated pits possibly represented ultrafast endocytosis intermediates, we localized a protein crucial for ultrafast endocytosis, Dyn1xA, in acute brain slices using stimulated emission depletion (STED) microscopy. The kinetics of ultrafast endocytosis are produced by pre-recruitment of endocytic proteins at the endocytic zone^32,33^. Dyn1xA molecules are therefore expected to localize next to the active zone of synapses at rest if those synapses can perform ultrafast endocytosis. Previous studies of Dyn1xA mainly relied on tagged protein constructs that were either overexpressed or genetically knocked-in to neurons^33,42^. To examine endogenous protein localizations in slices, we developed and knock-out validated a rabbit polyclonal antibody against a Dyn1xA specific C-terminal 20 amino acid region, which interacts with Endophilin A1 to mediate ultrafast endocytosis^32^ (Figure S4). Immunoblotting showed that the Dyn1xA antibody reacted with lysate from mouse neurons but not astrocytes (Figure S4A). Immunoblotting and fluorescence imaging showed the antibody was specific for the Dyn1xA splice isoform (Figure S4B,C). The selected amino acid region used as the antibody epitope is 100% identical between mouse and human (Figure S4E), and this antibody recognized Dyn1xA protein in mouse and human whole brain lysates (Figure S4D). Analysis of mouse cortical synapses showed the majority of punctate Dyn1xA antibody signals present near the presynapse of PSD95-positive excitatory synapses (Figure 4A-C; Figure S5 for individual signal traces and distances for each mouse; Figure S6A,B for cartoon representation of synapse orientation classification and example mouse slice STED overview; Figure S7 for STED analysis pipeline of side view synapses). Similar to cultured hippocampal neurons, Dyn1xA puncta were found directly at the active zone boundary and periactive zone (-50 nm to +50 nm) (Figure 4D,E), with nearly 47% of Dyn1xA signals localized within this region in excitatory cortical synapses (Figure 4F; see Figure S6C for STED analysis pipeline of top view synapses). These results suggest that Dyn1xA is present at the endocytic zone of cortical presynapses in acute mouse brain slices and advances the hypothesis that ultrafast endocytosis is responsible for millisecond synaptic vesicle recycling following a single stimulus in mice.

**Figure 4.**
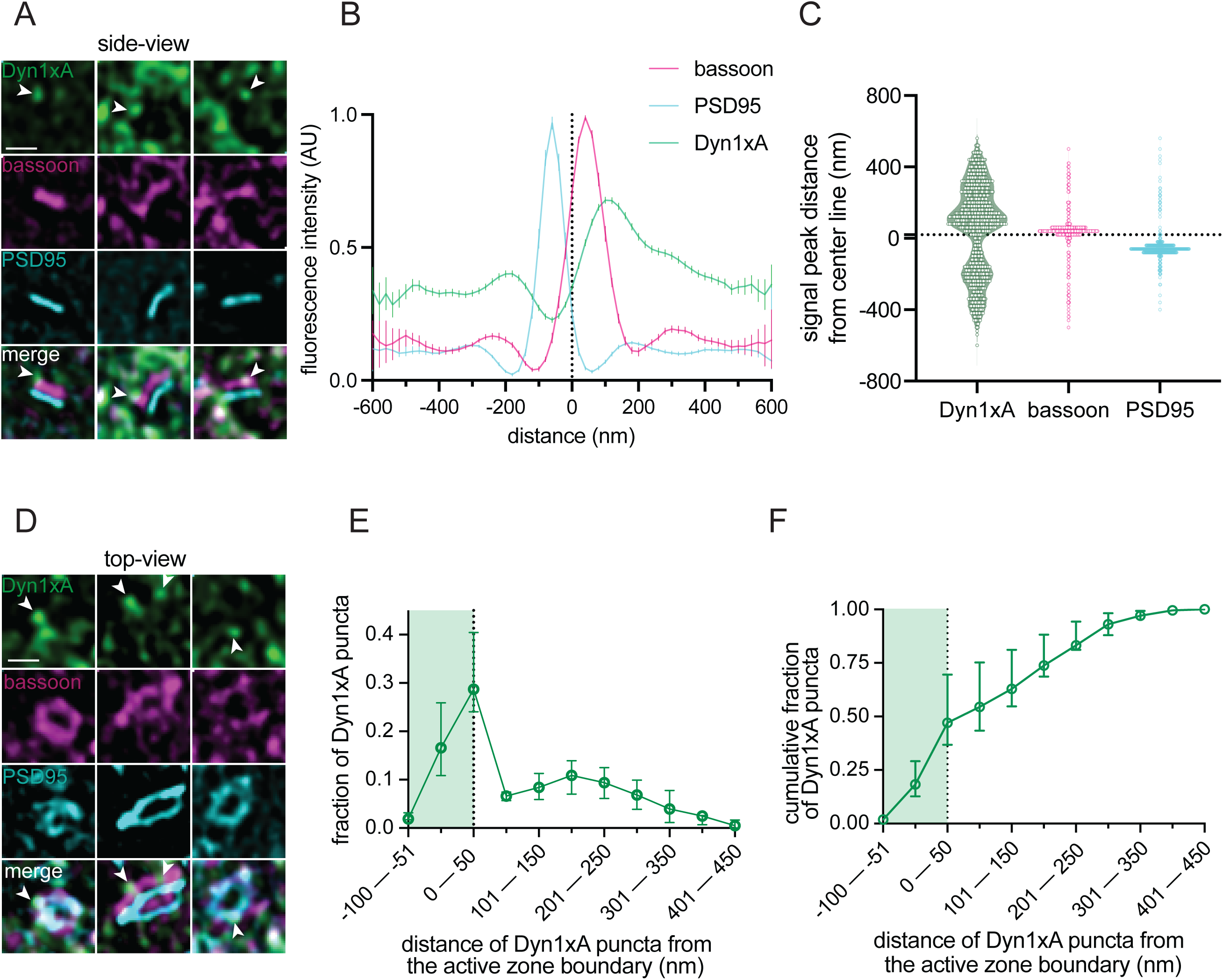
Dyn1xA clusters next to the active zone in mouse cortical synapses. (A) Example STED images of endogenous Dyn1xA localizations (white arrowheads) in side view synapses from cortical regions of acute mouse brain slices. Scale bar: 300 nm. Dyn1xA antibody confirmation in Figure S4. Individual traces provided in Figure S5. (B) Line scan profiles from side view synapses. The median and 95% confidence interval are shown for n=3 mice; see Data Table S1 for specific n values. Dotted line indicates center-line at x=0, representing the calculated center-line between the presynapse and postsynapse (see Figure S7 and Methods for analysis pipeline). (C) Violin plots of individual fluorescence signal peak distances from the center-line obtained from a band line scan across side view synapses. The median and 95% confidence interval are shown for n=3 mice, each dot represents a signal peak over a set threshold value >0.5. Dotted line indicates center-line at y=0. (D) Example STED images of endogenous Dyn1xA localizations (white arrowheads) in top view synapses from cortical regions of acute mouse brain slices. Scale bar: 300 nm. Dyn1xA antibody confirmation in Figure S4. More example STED images provided in Figure S6. (E) The distribution of Dyn1xA puncta relative to the active zone edge, defined by Bassoon, analyzed in top view synapse images. Shaded region indicates area inside the active zone. The median and 95% confidence interval are shown for n=3 mice; see Data Table S1 for specific n values. (F) Cumulative plot of data presented in (E).

### Ultrafast endocytosis in human cortical synapses

Having verified the zap-and-freeze approach in mouse cortical slices, we assessed its utility in human brain tissue. We received tissues from six epilepsy patients undergoing surgery (see Table S2 for de-identified patient information). These tissues originated from cortical areas and their removal was a required action in the overall surgery. We prepared 100 µm slices and performed our slightly modified NMDG recovery (see Methods for details, Figure S8A). To assess the viability and health of the slices, we performed electrophysiology for the first two cases before continuing. Recovered human pyramidal neurons (from 1 female and 1 male) showed typical action potential traces in response to depolarization, ability to respond to current steps (Figure S8B-G), and solid membrane properties with slight inter-individual variability as expected (Table S3). The recorded neurons also exhibited normal morphology, assessed with post-recording biocytin cell-fill staining (Figure S8H), indicating the tissues were viable and healthy. Tissue morphology was also largely preserved with the use of 15% PVP, and pearled axon morphologies were observed in human neocortical slices (Figure 5A i and ii).

**Figure 5.**
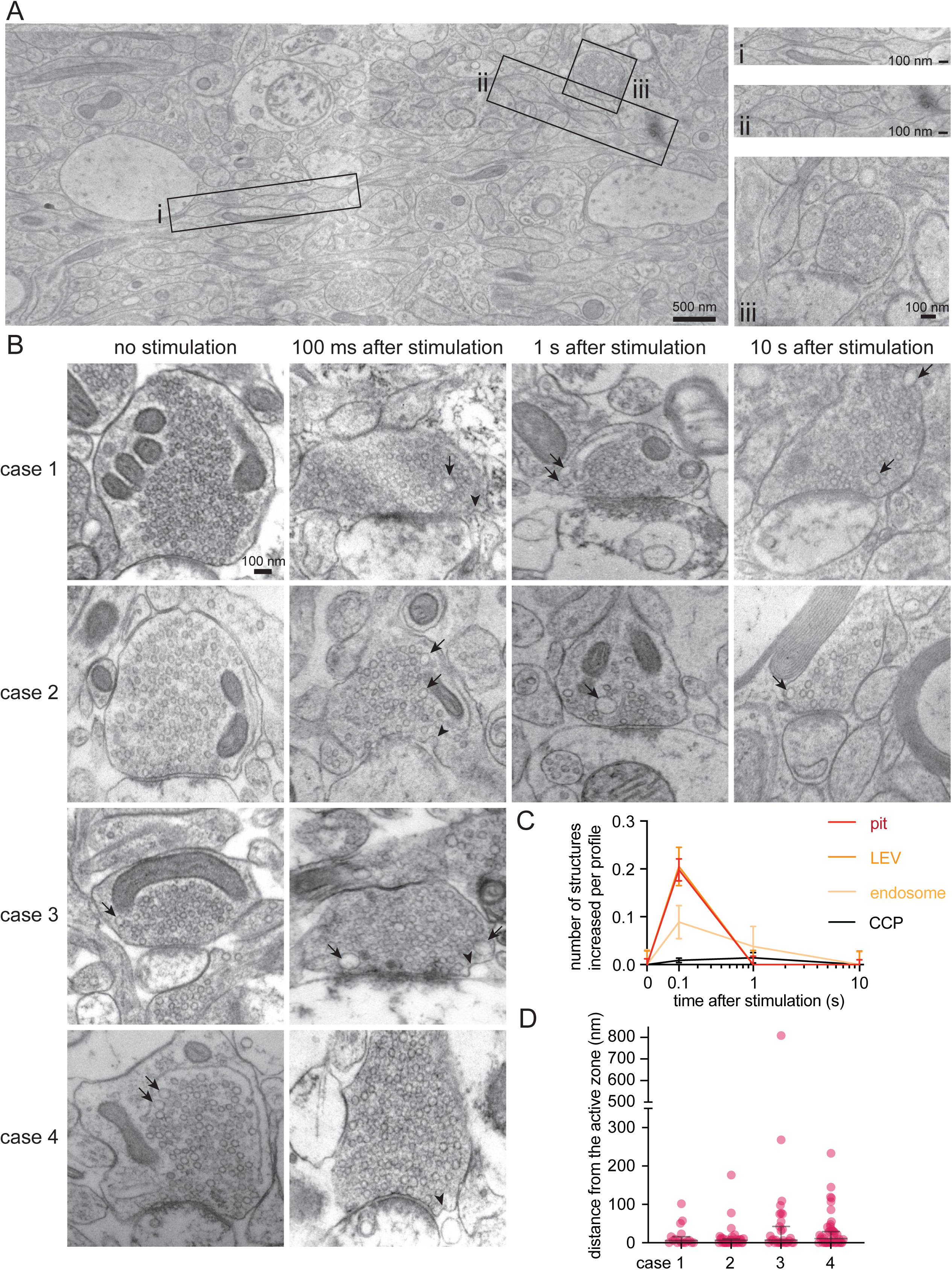
Ultrafast endocytosis in cortical synapses of acute human brain slices. (A) Low magnification overview of acute human neocortical slice visualized with zap-and-freeze EM (from a slice frozen 10 sec post-electric field stimulation). Axons with pearled morphologies are highlighted as panels (i and ii), along with an example of a synapse (iii). (B) Example electron micrographs showing uncoated endocytic pits (black arrowheads) and putative large endocytic vesicles (LEV) and endosomes (black arrows) at the indicated time points in acute brain slices from n=4 humans. Electrophysiological confirmation of human brain slice electrical viability in Figure S8. More example TEM images provided in Figure S9. (C) Plots showing the increase in number of each endocytic structure per synaptic profile after a single stimulus. Data are presented as mean ± SEM. CCP: clathrin-coated pits. See Data Table S1 for n values and detailed numbers for each time point. (D) Plot showing the distance distribution of putative uncoated endocytic pits from the edge of an active zone 100 ms post-stimulus in acute neocortical slices from n=4 humans. Data are presented as median ± 95% confidence interval. Each dot represents a pit.

As with the mice, we stimulated human slices from the first two cases (case 1 and 2, 1 female and 1 male) and froze them at 100 ms, 1 s, and 10 s. Unstimulated slices were used as controls. In both cases, uncoated pits appeared predominantly within 5-7 nm from an active zone, 100 ms after a single stimulus (Figure 5B-D, Figure S9A,B for more micrographs; median pit distances: 5.775 and 6.499 nm for case 1 and 2 respectively). Simultaneously, the number of large vesicles increased, indicating potential membrane internalization. Interestingly, the number of putative endosomes also peaked at 100 ms. To further probe the ultrafast endocytic pit formation timepoint, we froze two additional cases (case 3 and 4, 2 males) at 100 ms with the necessary unstimulated controls. Similarly, these two human cases exhibited uncoated pits within 6-10 nm from an active zone (Figure 5D; median pit distances: 6.600 and 10.76 nm for case 3 and 4 respectively; see Figure S9C,D for more micrographs), 100 ms after a single stimulus (Figure 5B,C). We then tested if these observed structures were activity or dynamin-dependent using tissue from 2 more human cases (case 5 and 6, 2 males), with unstimulated slices and non-drug treated slices from these same individuals as controls. Here uncoated pits formed 8-31 nm from an active zone (Figure S10C; median pit distances: 7.623 and 31.361 nm for case 5 and 6 respectively) 100 ms after a single stimulus in slices given no drugs. Similar to mouse cortical slices, pre-treatment of human cortical slices with TTX (1 µM, 10 min) abolished the prevalence of uncoated pits found at 100 ms post-stimulus (Figure 6A,B), while pre-treatment with Dynasore (80 µM, 2 min) trapped uncoated pits at the presynaptic plasma membrane 1 s post-stimulus (Figure 6A,C), suggesting that the formation of these uncoated pits is activity-dependent and Dynasore-sensitive in human cortical synapses.

**Figure 6.**
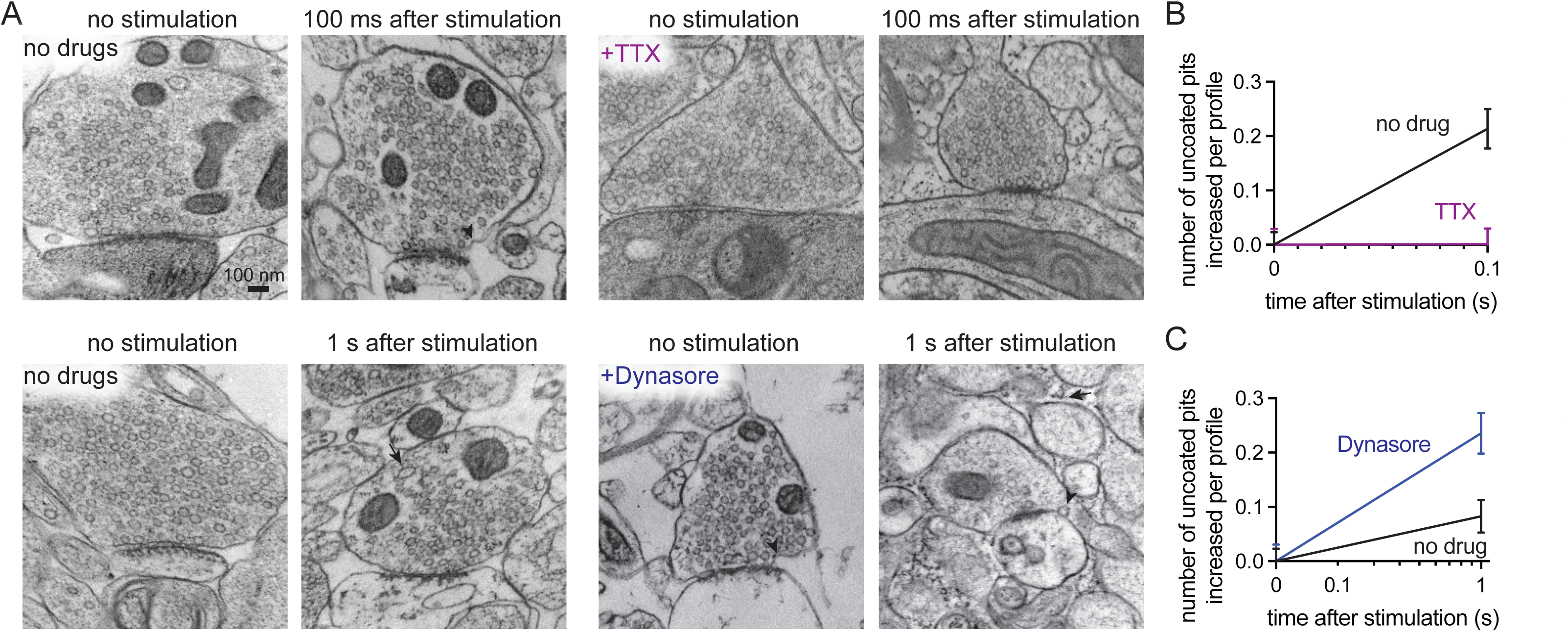
Uncoated pits in acute human cortical slices are activity-dependent and Dynasore-sensitive. (A) Example electron micrographs showing endocytic pits (black arrowheads) and putative large endocytic vesicles (LEV) at the indicated time points in acute brain slices from n=2 humans (case 5 and 6), given: no drugs, tetrodotoxin (TTX), or Dynasore. (B) Plot of no drug versus TTX treated slices, comparing the increase in number of uncoated pits per synaptic profile, 100 ms after a single stimulus. Data are presented as mean ± SEM. (C) Plot of no drug versus Dynasore treated slices, comparing the increase in number of uncoated pits per synaptic profile, 1 s after a single stimulus. Data are presented as mean ± SEM. See Data Table S1 for n values and detailed numbers for each time point.

Uncoated pits observed in the all six human cases forming at 100 ms post-stimulus had diameters ranging from 36-55 nm and neck widths ranging from 67-107 nm (Figure S10A, B; median pit diameters: 45.032, 46.494, 55.620, 44.076, 39.194, and 36.161 nm; median pit neck widths: 68.356, 90.091, 107.743, 80.773, 67.655, and 67.479 nm for case 1, 2, 3, 4, 5, and 6 respectively). Across all timepoints tested, human endosomes ranged in size from 6,019-10,386 nm^2^ (Figure S10D; median endosome sizes: 8,103.18, 6,380.86, 8,160.01, 10,386.30, 7117.98, and 6018.77 nm^2^ for case 1, 2, 3, 4, 5, and 6 respectively). Clathrin-coated pits were occasionally present but did not increase in numbers after stimulation at the timepoints examined (Figure 5C; see Figure S9E for micrographs of uncoated pits versus observed clathrin-coated pits from this dataset for comparison). No clear Ω-figures in the active zone, indicative of kiss-and-run like events, were observed in these human samples. These data suggest that synaptic vesicle endocytosis in human cortical synapses could be mediated by ultrafast endocytosis on a millisecond timescale following a single stimulus.

To interrogate a possible molecular pathway conservation, we localized Dyn1xA in acute human brain slices, prepared on the same day (see Methods). Notably, STED imaging across three cases (1 female and 2 males; note that we could not recover enough tissue from all cases) showed the majority of Dyn1xA puncta near the presynapse of PSD95-positive excitatory cortical synapses (Figure 7A,B; see Figure S11 for example human slice STED overview and additional side view synapses; Figure S12 for individual signal traces and distances for each human) located predominantly at the active zone boundary and periactive zone (-50 nm to +50 nm) (Figure 7C,D). Up to 46% of Dyn1xA signals localized within this region (Figure 7E; ∼42% for case 2, ∼46% for case 3, and ∼35% for case 4).

**Figure 7.**
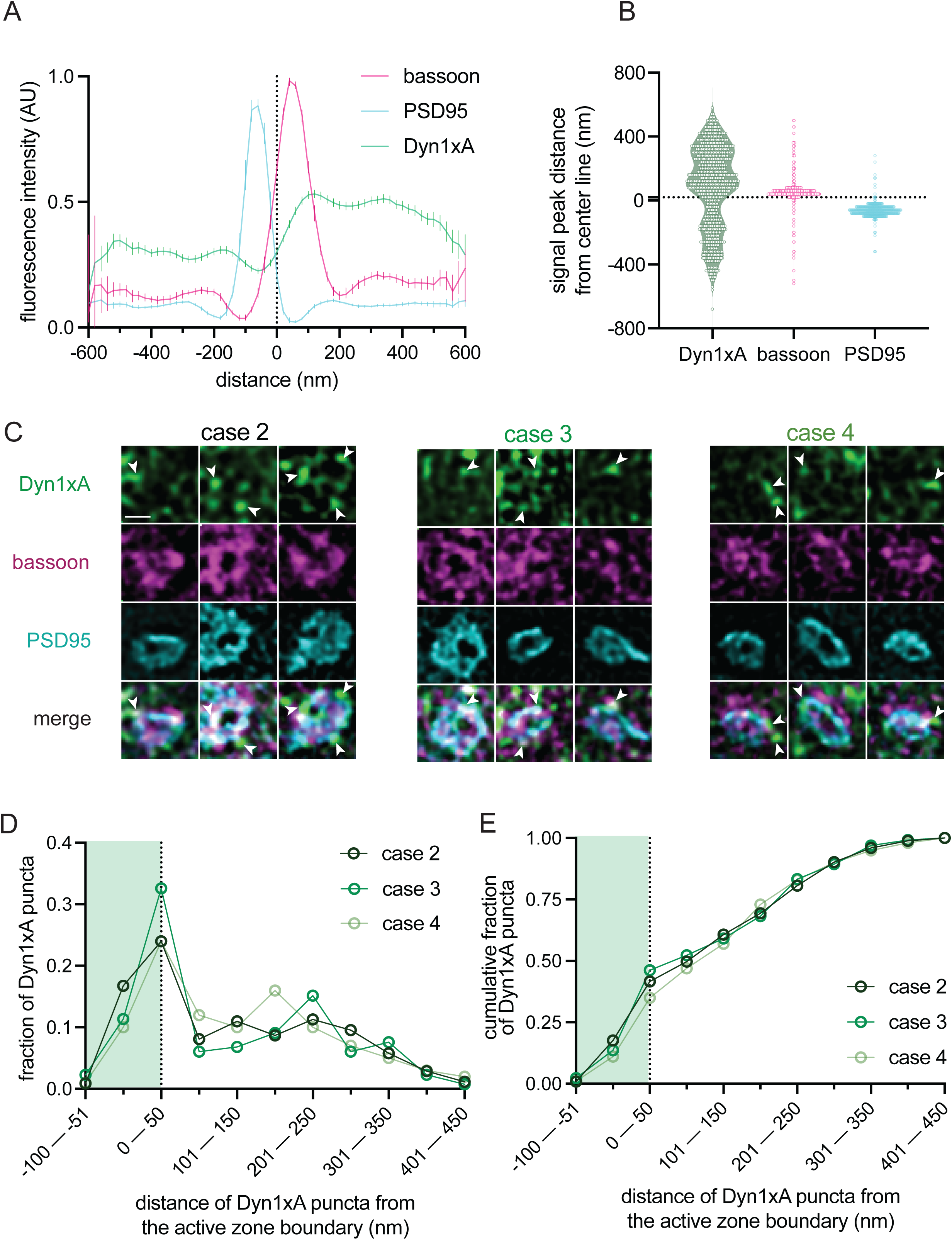
Dyn1xA clusters in human excitatory synapses. (A) Line scan profiles from side view synapses. The median and 95% confidence interval are shown for n=3 humans; see Data Table S1 for specific n values. Dotted line indicates center-line at x=0. (B) Violin plots of individual fluorescence signal peak distances from the center-line obtained from a band line scan across side view synapses. The median and 95% confidence interval are shown for n=3 humans, each dot represents a signal peak over a set threshold value >0.5. Dotted line indicates center-line at y=0. (C) Example STED images of endogenous Dyn1xA localizations (white arrowheads) in top view synapses from cortical regions of acute brain slices from n=3 humans. Scale bar: 300 nm. More example STED images provided in Figure S11. (D) The distribution of Dyn1xA puncta relative to the active zone edge, defined by Bassoon, analyzed in top view synapse images. Shaded region indicates area inside the active zone. See Data Table S1 for specific n values. (E) Cumulative plot of data presented in (D).

Altogether these data show that zap-and-freeze can be used to investigate the membrane dynamics of human brain tissues at millisecond and nanometer resolutions, and suggest that ultrafast endocytosis is possibly conserved in the human brain.

## Discussion

Here we present an approach to study membrane dynamics in acutely resected human brain tissue. The zap board for a high-pressure freezer induces neuronal activity in acute mouse cerebellar and neocortical slices, as detected by calcium imaging. Ultrastructural analysis of mouse and human cortical synapses showcase similar uncoated pits forming ≤31 nm from the active zone 100 ms post-stimulus. Localizations of Dyn1xA near the endocytic zone of mouse and human cortical synapses, alongside micrographs illustrating observed structures are activity-dependent and Dynasore-sensitive, suggest that these uncoated pits likely represent ultrafast endocytosis intermediates.

### Strengths and shortcomings of the method

The presented method has both advantages and disadvantages. A major advantage is that no exogenous proteins like channelrhodopsin are necessary to activate neurons. Therefore, we can attempt to study synaptic transmission in a more ‘native’, *ex vivo* context and in organisms or tissues not easily genetically manipulated. Patient tissues also maintain cell diversity, interactions, and cytoarchitectures that stem cell *in vitro* systems cannot provide^43^ and thus, may reveal cell-autonomous mechanisms occurring in neurons and non-cell autonomous events involving surrounding glia^44^. Zap-and-freeze may prove useful in examining post-mortem or surgical resections from patients affected by disease (provided short post-mortem and tissue acquisition time intervals). Many neurological and neurodegenerative diseases present synaptic dysfunction^45^ and endolysosomal defects^46^. Visualizing synaptic dynamics in human tissue is likely to yield more directly translatable results for potential treatment developments and may facilitate the validation of cell biology mechanisms identified in animal and cell models.

It is worth noting that this approach has limitations. We tested neocortical slices from male (n=5) and female (n=1) epilepsy patients spanning an approximate age range from 20 to 40 years old. The parameters we measured seemed unaffected by sex assigned at birth, age, or possible past brain traumas. However, we cannot dismiss the fact that this is likely due to the limited number of independent specimens. We also do not have access to genetic or other private patient information, which may affect the interpretation of results. From a technological standpoint, our theoretical calculations suggest that it would take an additional 3-4 ms for the core of brain slices to reach 0 °C ^19^, leaving an ambiguity in the timing of the actual freeze relative to stimulation. This issue would not affect the method’s ability to capture synaptic vesicle endocytosis but presents a possible challenge for investigating exocytosis mechanisms that occur more quickly. Our use of 2D imaging is also probably excluding subtle, interesting information about intra-tissue connections, though we expect this technique can be easily adapted for use with existing 3D EM imaging. Furthermore, unlike optogenetic stimulation, most axons parallel to the electric field are likely stimulated, making it more difficult to study circuit-specific questions. Finally, as with other time-resolved electron microscopy approaches, we cannot follow the membrane dynamics of the exact same synapses over time like what can be accomplished via live-cell imaging. Thus, our conclusions are drawn from the most frequent events observed at each time point snapshot; it is therefore possible to miss less frequent events like kiss-and-run and clathrin-mediated endocytosis. To compensate for this restriction, future development of live super-resolution imaging of endocytic and fusion pore dynamics^47^ is warranted in brain slices^48^. Nonetheless, the potential benefits of zap-and-freeze as a technique outweigh these challenges.

### Ultrafast endocytosis in human brain

Mechanisms of synaptic vesicle recycling have been debated over the last 50 years. In the 1970’s, Heuser and Reese proposed clathrin-mediated endocytosis as the major pathway for synaptic vesicle recycling after intense stimulation of frog neuromuscular junctions^49^. Around the same time, Ceccarelli and colleagues proposed another mechanism^39^, named kiss-and-run^50^. Over the years, evidence for both mechanisms has accumulated^51^. Since the early observation of fast compensatory endocytosis in bipolar cells of fish^52^, we have presented evidence over the last decade for clathrin-independent, ultrafast endocytosis occurring in both *Caenorhabditis elegans* neuromuscular junctions and mouse hippocampal synapses^12,30,31,33–35,42^. Similarly, several groups have implicated the existence of ultrafast mechanisms at play in mammalian central synapses^53–56^. Our data here add to this hypothesis.

Taking into account that fluid phase markers are not used in this study, we cannot exclude the possibility that the uncoated pits appearing at 100 ms could represent exocytic intermediates of large vesicles, which may have formed through fusion of two or more synaptic vesicles (i.e. ‘compound fusion’)^57^. In chromaffin cells, such compound fusion occurs sequentially: one vesicle fuses with the plasma membrane, and another vesicle fuses directly onto the first as they are incorporated into the plasma membrane^58^. Sequential compound fusion is a reasonable mechanism for chromaffin cells as fusion of granules with diameters ∼200 nm takes seconds to occur—providing enough time for t-SNARE complexes to theoretically diffuse into the fusing membrane. However, synaptic vesicles are ∼40 nm in diameter, and exocytosis completes within 10 ms^19^, making it unclear how t-SNAREs could be available on fusing vesicles in such a short time frame. Nonetheless, further studies are warranted to investigate the pervasiveness of compound fusion in the human brain.

Using zap-and-freeze EM we were able to conduct a brief comparison of activated mouse and human cortical synapses from intact tissue. Our data indicate that the timing of synaptic vesicle recycling is relatively similar between mouse and human cortical synapses. Yet, it seems as though the uncoated pits forming at 100 ms are slightly larger in diameter with wider necks in humans compared to mice. Our data also indicate that in general human endosomes are larger than those found in mice. From this initial study, we cannot determine if these differences are species-specific or possibly due to these human tissues originating from epileptic brains.

Though we used a single stimulus to probe synaptic vesicle endocytosis, this stimulus is likely relevant given that the firing rate of some human cortical neurons can be very low based on the cognitive task being performed (<1 Hz)^59^. More importantly, the molecular mechanism may be conserved in human cortical synapses. As in cultured mouse hippocampal neurons, the endocytic protein Dyn1xA appears to be pre-localized to the endocytic zone, potentially able to accelerate endocytosis at human synapses. Though mechanistic investigations of lab-based organisms will undoubtedly contribute to our understanding of synaptic transmission, direct examination of human tissues continues to provide key information.

## Acknowledgments

We would like to thank Kristen Harris for providing invaluable advice on the recovery timing needed for human brain slices at the beginning of this work. We also thank Barbara Smith, Mike Delanoy, LaToya Roker, Loza Lee, and Scot Kuo at the Johns Hopkins Microscopy Facility for technical and administrative assistance in electron microscopy; Aleksandr Smirnov and Susan McTeer at the Johns Hopkins Neuroscience Imaging Center for technical and administrative assistance in STED microscopy; Abbas Shirinifard at the St. Jude Neuroimaging Analysis Laboratory for technical assistance in machine-learning approaches and Python code generation. We especially thank all patients who lend their tissue to research—we would find nothing without them and their extraordinary contributions.

C.R.E. was supported by the Howard Hughes Medical Institute (HHMI) Gilliam Fellowship for Advanced Study (GT14961). Y.I. was supported by the Kazato Foundation and American Lebanese Syrian Associated Charities (ALSAC). P.F.W. was supported by the National Institutes of Health (U19 AG072643). K.L. was supported by Marine Biological Laboratory Whitman Fellowship, a Young Investigator Grant from the Medical Faculty, Leipzig University, and a fund (‘ASAP’) from the Roland Ernst Stiftung, Germany. S.W. was supported by funds from the Johns Hopkins University School of Medicine, Marine Biological Laboratory Whitman Fellowship, Chan-Zuckerberg Initiative Collaborative Pair Grant, Chan-Zuckerberg Initiative Supplement Award, Brain Research Foundation Scientific Innovation Award, Helis Foundation award, the Kleberg Foundation grant, and the National Institutes of Health (1DP2 NS111133-01, 1R01 NS105810-01A1, R35 NS132153) awarded to S.W. S.W. is an Alfred P. Sloan fellow, a McKnight Foundation Scholar, a Klingenstein and Simons Foundation scholar, and a Vallee Foundation Scholar. The EM ICE high-pressure freezer was purchased partly with funds from an equipment grant from the National Institutes of Health (S10RR026445) awarded to S.C. Kuo. This manuscript is the result of funding in whole or in part by the National Institutes of Health (NIH). It is subject to the NIH Public Access Policy. Through acceptance of this federal funding, NIH has been given a right to make this manuscript publicly available in PubMed Central upon the Official Date of Publication, as defined by NIH.

## Author contributions

**Chelsy Eddings:** conceptualization, mouse and human cortical brain slicing and recovery, TTX and Dynasore testing of mouse and human brain slices, zap-and-freeze EM, TEM image acquisition and analysis, immunohistochemistry, STED imaging and analysis, writing—original draft and figure construction

**Minghua Fan:** electrophysiology and analysis, biocytin imaging

**Yuuta Imoto:** STED image analysis code construction, Dyn1xA antibody design and validation

**Kie Itoh:** Dyn1xA antibody design and validation

**Xiomara McDonald:** TEM image analysis

**Jens Eilers:** conceptualization of zap board testing and 2-photon calcium imaging with mouse cerebellum in Figure 1 and S1

**William Anderson:** human tissue surgical acquisition

**Kristina Lippmann:** conceptualization of zap board testing with mouse cerebellum and cortex, mouse cerebellar and cortical brain slicing and recovery, 2-photon calcium imaging, zap board testing

**Paul Worley:** supervision of human cortex electrophysiology

**David Nauen:** conceptualization, human tissue pathology assessment

**Shigeki Watanabe:** conceptualization, TEM image analysis, writing—original draft and figure construction

*all authors edited the manuscript

## STAR Methods

### Lead contact

Further information and requests for resources and reagents should be directed to and will be fulfilled by the lead contact, Shigeki Watanabe.

### Data and code availability

All original code and data are available from the lead contact upon request. Any additional information required to reanalyze the data reported in this paper is available from the lead contact upon request. The macros and Matlab scripts for EM image analysis are available at https://github.com/shigekiwatanabe/SynapsEM). Additional procedures are described in Watanabe et al. (2020)^60^. STED image analysis codes are available at (https://github.com/imotolab-neuroem/STED_image_analysis_package_public_v1.4/tree/main).

### Mice

All procedures involving mice were approved by the Johns Hopkins Animal Care and Use Committee and the regional authority of the state of Saxony, Germany (T09/16, T10/20), following the guidelines of the National Institutes of Health and the European Communities Council. Both males and females were used in this study. All animals were kept on a 12 hour light/dark cycle and provided access to unlimited food and water. Wild-type mice used for slice experiments were obtained from Taconic (B6NTAC), kept at Johns Hopkins, and from in-house breeding (C57BL/6N) from the medical experimental center at Leipzig University. *Dnm1* KO mice^61^ used for Dyn1xA antibody validation were obtained from Dr. Pietro De Camilli.

### Human tissue

Patient samples were collected under a protocol by the Institutional Review Board (IRB) at Johns Hopkins Hospital from neurosurgical resections performed for neurologic disease, generally intractable epilepsy. Because the tissue resections occurred regardless of any research use, the IRB determined that waiver of consent was appropriate. After gross assessment by a neuropathologist, a small portion of diagnostically unnecessary neocortical tissue was de-identified and placed in room temperature NMDG containing artificial cerebrospinal fluid (aCSF) while transported to the lab—recipe described below.

### Acute brain slice preparation

Acute mouse cerebellar and cortical slices for 2-photon imaging on the zap board were prepared from 21- to 26-day-old C57BL/6N mice of either sex (n=8), as described previously^25^. In short, under deep isoflurane anesthesia mice were decapitated. The skull was immediately opened, and the cerebellar vermis or prefrontal neocortex was rapidly removed and placed in ice-cold aCSF_2 (in mM: 125 NaCl, 2.5 KCl, 2 CaCl_2_, 1 MgCl_2_, 1.25 NaH_2_PO4, 26 NaHCO3, 20 glucose, bubbled with 95% O_2_ and 5% CO_2_, pH 7.4 at room temperature). Using a vibratome (Leica VT1200S), 100 µm horizontal slices of the cerebellar vermis or sagittal slices of the prefrontal cortex were sectioned and incubated in aCSF_2 at 35°C for 30-40 min before they were stored at room temperature until usage.

For zap-and-freeze and STED experiments, mouse and human acute cortical slices were cut to 100 µm on a vibratome (Leica VT1200S) in room temperature NMDG aCSF (in mM: 92 NMDG, 2.5 KCl, 1.25 NaH_2_PO4, 30 NaHCO_3_, 20 HEPES, 25 glucose, 2 thiourea, 5 Na-ascorbate, 3 Na-pyruvate, 10 NAC, 0.5 CaCl_2_·2H_2_O, and 10 MgSO_4_·7H_2_O). Brain slices were then transferred into a custom-built recovery chamber filled with continuously carbogenated NMDG solution at 37 °C and allowed to undergo an initial 12 min recovery. After 12 min, slices were moved into a new recovery chamber filled with continuously carbogenated HEPES holding aCSF (in mM: 92 NaCl, 2.5 KCl, 1.25 NaH_2_PO_4_, 30 NaHCO_3_, 20 HEPES, 25 glucose, 2 thiourea, 5 Na-ascorbate, 3 Na-pyruvate, 2 CaCl_2_·2H_2_O, and 2 MgSO_4_·7H_2_O) and allowed to undergo a second recovery for at least 4 hr at 37 °C. Slices were recovered to ensure electrical viability and responsiveness to later electric field stimulation—also to repair cut site damage to maximize the amount of maintained tissue morphology while imaging. All aCSF solutions were titrated to 7.3-7.4 pH, left at room temperature to avoid potential tissue shrinkage-expansion cycles that can affect tissue ultrastructure, and bubbled with carbogen gas (95% O_2_, 5% CO_2_) before use. Note that we did not perform the Na+ spike-in procedure associated with the original NMDG recovery method^29^ to prevent possible tissue overexcitation.

### 2-photon calcium imaging on the zap board

For calcium imaging, cerebellar or cortical slices were incubated for 20 min with 4 µmol Fura-2, AM (solved in Pluronic F-127 20 % solution in DMSO) in a submerged recording chamber filled with aCSF_2. To ensure sufficient incubation, circulation of carbogenated aCSF_2 was stopped, and carbogen supply was provided above the fluid with a sinter filter and a cardboard box on top to create an oxygen-rich environment. After incubation, excess dye was washed out for another 20 min with carbogenated, circulating aCSF_2. Slices were then transferred with a spatula and paintbrush to the zap board and assembled in aCSF_2 between two sapphire discs, separated by a 100 µm Mylar spacer. A rubber O-ring was placed on top of the assembly to imitate high-pressure freezing experiments. Electrical TTL-driven inputs to the zap board were driven by an EPC-10 USB patch clamp amplifier and Patchmaster software (HEKA Elektronik), and signal outputs digitized. Voltage for triggering the LED in the optocoupler was set to 2.55 V for slice experiments. Calcium imaging was performed with a 2-photon laser scanning microscope (BX61WI microscope and Fluoview 10 M scanner, Olympus), using a 50 mm, f1.2 camera objective (Zuiko, Auto-S OM-system, Olympus) and a mode-locked Ti:sapphire laser (Mai Tai DeepSee, Spectra-Physics, set to a center wavelength of 800 nm), controlled by Olympus Fluoview ASW software (version 04.01). Line scans were performed to test stimulation paradigms, at room temperature. The calcium signal after stimulation was calculated over background (dF/F_0_). Minimum Fura-2, AM signal was revealed by exponential fits. Data was normalized to the minimum for comparison with other slices. For cortical traces, the area of the calcium signal was calculated from the first stimulus (70 ms) until the end of the scan (900 ms), measuring between the baseline value (at y = 1, dotted line) and the trace.

### Zap-and-freeze experiments

For zap-and-freeze of acute mouse and human brain slices, tissue was always tested the same day as collection. Slices were transported from recovery chambers to the high-pressure freezer in a petri-dish filled with room temperature aCSF (in mM: 125 NaCl, 2.5 KCl, 1.25 NaH_2_PO_4_, 25 NaHCO_3_, 10 glucose, 2 CaCl_2_·2H_2_O, and 2 MgSO_4_·7H_2_O; 7.3-7.4 pH). Slices were then trimmed into smaller pieces ∼6 mm by hand using a razor blade. After trimming, slices were placed into freezing medium containing pre-warmed aCSF, supplemented with NBQX (3 mM), Bicuculline (3 mM), and 15% polyvinylpyrrolidone (PVP) as a cryoprotectant—slices were kept in this solution for less than 2 min as the freezing apparatus was assembled. 15% PVP was chosen and assembled in a specimen sandwich based on a method previously published for optogenetic stimulation of mouse brain slices using the high-pressure freezer^13^. The table and sample chamber of the high-pressure freezer were kept at 37 °C to ensure the physiological temperature of slices were maintained during experiments^62^. Unstimulated controls for each experiment were always frozen on the same day and originated from the same mouse or patient. The device was set such that samples were frozen at 0.1, 1 or 10 sec after the stimulus initiation. Cortical columns were always visually aligned perpendicular to the direction of the electric field to maintain consistency across samples (this ensured cortical layers were in parallel with the electric field).

### Pharmacological experiments

To test the potential activity-dependence of uncoated pit formation: after recovery, acute slices were placed in room temperature, continuously carbogenated aCSF supplemented with 1 µM tetrodotoxin (TTX) for 10 min before being placed into pre-warmed freezing medium and immediately undergoing zap-and-freeze. To test the potential dynamin-dependence of uncoated pits: after recovery, acute slices were placed in a petri-dish containing room temperature aCSF supplemented with 80 µM Dynasore for 2 min before being placed into pre-warmed freezing medium and immediately undergoing zap-and-freeze.

### Freeze substitution

Frozen samples were transferred under liquid nitrogen to an automated freeze substitution unit held at -90 °C (EM AFS2, Leica Microsystems). Using chilled tweezers, samples were moved into acetone to help disassemble the freezing apparatus. Sapphire disks with brain slices were then quickly moved into sample carriers containing 1% glutaraldehyde (GA) and 0.1% tannic acid (TA) in anhydrous acetone. The freeze substitution program was as follows: -90 °C for at least 36 hrs with samples in 1% GA, 0.1% TA, samples were then washed five times with pre-chilled acetone (30 min each), after washing the fixative solution was replaced with pre-chilled 2% OsO_4_ in acetone and the program was allowed to continue; -90 °C for 11 hrs with samples in 2% OsO_4_, +5°C per hour to -20 °C, -20 °C for 12 hrs, +10 °C per hour to +4 °C, hold at +4 °C.

### Sample preparation for electron microscopy

After freeze substitution, samples removed from the AFS were washed six times with acetone (10 min each) and incubated with increasing levels of plastic (100% epon-araldite diluted with acetone: 30% for 2 hrs, 70% for 3 hrs, and 90% overnight at +4 °C). After plastic infiltration, samples were embedded in 100% epon-araldite resin (Araldite 4.4 g, Eponate 12 Resin 6.2 g, Dodecenyl Succinic Anhydride (DDSA) 12.2 g, and Benzyldimethylamine (BDMA) 0.8 ml) and cured for 48 hrs in a 60 °C oven. Serial 40-nm sections were then cut using an ultramicrotome (EM UC7, Leica microsystems) and collected onto 0.7% pioloform-coated single-slot copper grids. Sections were stained with 2.5% uranyl acetate in a 50-50 methanol-water solution.

### Transmission electron microscopy

Images of synapses were acquired on a Hitachi 7600 transmission electron microscope at 80 kV and 80,000x magnification, with a AMT XR80 high-resolution (16-bit) 8-megapixel CCD camera, using AMT Image capture engine V602 (AMTV602) software. Samples were blinded and given random names before imaging. At least 98-100 images were obtained per experimental timepoint. Low magnification images of tissues were obtained at 30,000x (mouse) and 40,000x (human) magnification and stitched together using the TrakEM2 plugin for Fiji^63^, https://github.com/trakem2, with final contrast adjustments made in Adobe Photoshop (v21.2.1).

### Electron micrograph image analysis

Electron micrographs were manually analyzed using the published SynapsEM protocol^60^. Images from one experimental replicate were pooled into a single folder, randomized, and blinded using Matlab scripts. Synapses not containing a prominent postsynaptic density or those with poor morphology were manually excluded from analysis after this blinded randomization. Using custom Fiji macros, membrane and organelle features were annotated and exported as text files. Those text files were again imported into Matlab where the number and locations of the annotated features were calculated. For pit distribution from the active zone, the distance from the nearest edge of pits to the active zone was calculated. To minimize bias and error all annotated, randomized images were thoroughly checked and edited by at least one other member of the lab. Representative electron micrographs were adjusted in brightness and contrast to different degrees, rotated and cropped in Adobe Photoshop (v21.2.1) or Illustrator (v24.2.3). All Fiji macros and Matlab scripts are publicly available at (https://github.com/shigekiwatanabe/SynapsEM).

### Antibody generation

To generate the Dyn1xA antibody, the C-terminal Dyn1xA-specific residues [846-864, Cys-RSGQASPSRPESPRPPFDL] were custom synthesized and used for rabbit immunization (Pacific Immunology). The antibody was affinity purified using the antigen peptides.

### Immunohistochemistry

For immunohistochemistry, adult mice were anesthetized with isoflurane, transcardially perfused with 4% paraformaldehyde (PFA) and decapitated, with their whole brain dissected into PBS. Whole brains were embedded in O.C.T. compound (Tissue-Tek), frozen on dry ice, and sectioned in a coronal orientation at 40 µm on a cryostat (Leica CM 3050S). Human brain slices were collected from the tissue remaining after zap-and-freeze experiments were completed—slices were kept at 100 µm and fixed in 4% PFA. Staining was performed on free-floating sections according to the protocol described in Kruzich, E, et al^64^. Slices were permeabilized and blocked with 10% Normal Goat Serum (NGS) and 1% Triton X-100 in PBS for 3 hrs at room temperature. Primary antibodies diluted in 10% NGS and 0.025% Triton X-100 in PBS were added to slices for 48 hrs at +4 °C. Primary antibodies included: FluoTag-X2 anti-PSD95 (1:100), Bassoon (1:100), and Dynamin1xA (1:100). After removal of primary antibodies, slices were washed four times with 0.025% Triton X-100 in PBS, 15 min each wash. Secondary antibodies were diluted in PBS and incubated with slices for 48 hrs at +4 °C. Secondary antibodies included: goat anti-rabbit Alexa Fluor 594 (1:100) and STAR 460L goat anti-mouse IgG (1:100). Slices were washed four times with PBS, then washed once with Milli-Q water before being mounted onto plain microscope slides (Globe Scientific) using a paintbrush. Tissue was allowed to dry until transparent under foil. ProLong Diamond Antifade Mountant was added directly onto slices, with a coverslip (18 mm diameter 1.5H thick coverslips, Neuvitro Corporation) placed on top. Samples were left to dry at least overnight before STED imaging.

### Stimulated emission depletion (STED) imaging

2D STED images were acquired on an Aberrior FACILITY line microscope using a 60x oil objective lens (NA = 1.42). The excitation wavelengths were set as: 640 nm, 561 nm, and 485 nm for imaging Atto-643, Alexa-594, and STAR460L labeled target respectively. Imaging was performed at 20 nm pixel size, 0.61 AU pinhole, dwell time 5 µs. The STED beam was set at 775 nm with power of 10-15% used.

### STED image deconvolution, segmentation and analysis

#### i. *Image deconvolution*

1) STED images were exported from the microscope in .obf format using LiGHTBOX software (Abberior). Initial image processing was performed using a custom made MATLAB code package (Imoto et al. 2024^32^).
2) Images were extracted from .obf files and converted to .tif format as unsigned 16-bit integers. The extracted images were normalized based on the minimum and maximum intensity values within each image and then blurred with a Gaussian filter with 1.2 pixel radius to reduce the Poisson noise.
3) Subsequently, the overview tissue images were deconvoluted twice using the two-step blinded deconvolution method. The initial point spread function (PSF) input was measured from the unspecific antibody signals of STAR 460L, Alexa 594, or Atto 643 in the STED images. The second PSF (enhanced PSF) input was chosen by the user as the returned PSF from the initial run of blind deconvolution^65^. The enhanced PSF was used to deconvolute the STED images to be analyzed. Each time 10 iterations were performed.

#### ii. *Segmentation*

1) Series of the deconvoluted STED images were loaded to the segmentation script utilizing MIJ: Running ImageJ and Fiji within Matlab (Sage 2017, MATLAB Central File Exchange).
2) All presynaptic boutons in each deconvoluted image were indiscriminately selected within 45×45-pixel regions of interest (ROIs) based on the Bassoon and Dyn1xA signals.
3) Top view and side view presynapses were sorted via script with supervision based on Bassoon shape.

#### iii. *Side view synapse analysis*

Distance relative to the presynapse or postsynapse analysis was performed on side view images only. All side view synapse image processing was automated via a Python workflow integrating machine-learning based masking and geometric alignment. Analyses were run in Python 3.13.5 (Anaconda distribution) using Jupyter within Visual Studio Code (VS Code). For machine-learning, ilastik 1.4.0 (https://www.ilastik.org/) was invoked headlessly via the command line. Portions of this code were generated with assistance from GitHub Copilot (Python extension in VS Code) and the Anthropic Claude Sonnet 4.0 model.

1) For each mouse or human dataset, the three fluorescent channels were assigned as CH1= Bassoon, CH2= PSD95, and CH3= Dyn1xA. Pixel Classification projects for Bassoon and PSD95 were trained for each dataset using 5–7 randomly selected images (selected within the code) and applied Color/Intensity and Edge features at Gaussian scales σ = 0.3, 0.7, 1.0, 1.6, 3.5, 5.0, and 10.0. ilastik segmentations were read as label images and binarized to foreground=1/background=0. Foreground polarity was enforced so that the object class was mapped to 1.
2) For each image, the largest external contour of each channel’s ilastik mask (Bassoon and PSD95) was fit with an ellipse (OpenCV fitEllipse). The major-axis angle was defined as the ellipse’s long axis (angles normalized to ensure the longer diameter was always used). A ‘center-line’ (“AZ–PSD center line”) was then computed as the line parallel to both major-axes and passing through the midpoint between the two ellipse centers. When the two axes were nearly parallel (≤ 30° difference), their angles were averaged with wrap-around handling; otherwise, a vector (atan2-based) average was used. The center-line length was set to the mean of the two major-axis lengths.
3) Each image was then rotated to align this center-line vertically (rotation angle = −(90° − center-line angle)). Image orientation was always standardized such that Bassoon signals lay on the right of the center-line and PSD95 on the left of the center-line, by conditionally applying an additional 180° flip based on the Bassoon/PSD95 centroids relative to the center-line perpendicular. Rotational Not a Number values (NaNs) introduced by fifth-order spline interpolation (scipy.ndimage.rotate, order= 5) were replaced by the mean intensity of the corresponding channel.
4) For visualization, channel display ranges were set to the 1^st^-99th percentiles per image. For analysis, the band-scan profiles were min–max normalized to [0,1] per channel. A vertical band line scan was performed on the rotated images using a fixed band width of 42 pixels centered on the image midline (1 pixel = 20 nm). For each x-position, the mean intensity within the band-scan was computed, yielding one line scan profile per channel. The x-axis was then recentered such that 0 always corresponded to the center-line (left negative, right positive).
5) Dyn1xA signal peaks were called on the normalized CH3 line profile using scipy.signal.find_peaks, and signal peaks with amplitude > 0.5 were retained as real signals. For each image, all X-Y values underlying the band-scan line profiles (distance in pixels from the center line and normalized intensities for CH1-CH3) and the Dyn1xA peak positions were written as .csv files. To ensure transparency and reproducibility, intermediate composite figures (raw input channels, ilastik masks with fitted ellipse major axes, the computed parallel center-line, rotated channels, and band-scan overlays) were each saved per image as PNG files in a dedicated result/ directory. The corresponding ilastik-trained .ilp project files were also saved.
6) The numbers in the resulting .csv files were used to construct line scan profiles and violin plots in Prism.

#### iv. *Top view synapse analysis*

Distance distribution analysis was performed on top view images only, since Dyn1xA puncta can appear to be in the middle of synapses when seen in side view. All top view synapse imag processing was performed using a custom made MATLAB code package (Imoto et al. 2024^32^).

1) The boundary of the active zone was identified as the contour that represents half of the intensity of each local maxima in the Bassoon channel. The Dyn1xA puncta were picked by calculating pixels of local maxima.
2) Distances between Dyn1xA puncta and the active zone boundary were automatically calculated correspondingly. For this distance measurement, first, MATLAB distance2curve function (John D’Errico 2024, MATLAB Central File Exchange) calculated the distance between the local maxima pixel and all the points on the contour of the active zone or Dyn1xA cluster boundary. Next, the minimum distance for each local maxima pixel was selected. Signals crossing the ROIs and the Bassoon signals outside of the stained neurons were excluded from the analysis. The MATLAB scripts are available from github (https://github.com/imotolab-neuroem/STED_image_analysis_package_public_v1.4) or by request.
3) Values in the resulting MATLAB files were used to construct plots in Prism.

### Electrophysiological recordings, analysis and biocytin imaging

Human brain slices were transferred to a submerged recording chamber, perfused and maintained at 34°C in solution containing (in mM): 125 NaCl, 2.5 KCl, 1.25 NaH_2_PO4, 25 NaHCO_3_, 2 CaCl_2_, 2 MgCl_2_, 10 glucose, saturated with 95% O_2_, 5% CO_2_. Whole-cell recordings from human cortical pyramidal neurons were visualized under an upright Zeiss microscope with infrared optics. Recording pipettes had resistances of 3-5 MΩ when filled with an internal solution containing (in mM): 120 K-gluconate, 15 KCl, 10 HEPES, 2 MgCl_2_, 0.2 EGTA, 4 Na_2_ATP, 0.3 Na_3_GTP, and 14 Tris-phosphocreatine (pH 7.3). Whole-cell current-clamp recordings were made with MultiClamp 700B and 1440A digitizer (Molecular Devices). Data were acquired and analyzed using pClamp 10.4 (Molecular Devices). Series resistance was monitored throughout the recording and controlled below 20 MΩ. Data was discarded when the series resistance varied by ≥ 20%. Recorded neurons were filled with 0.2% (w/v) biocytin. Then, the slices containing biocytin-filled neurons cross-linked with 4% PFA and stained with Streptavidin Alexa Fluor 555 conjugate (1:1,000). Fluorescent images were obtained using a confocal microscope (Zeiss LSM 900).

### Expression constructs

To generate lentiviral expression constructs for Dyn1xA-FLAG and Dyn1xB-FLAG, f(syn)NLS-RFP-P2A-Dyn1xA and f(syn)NLS-RFP-P2A-Dyn1xB (Imoto et al. 2022^33^) were amplified and linearized using an In-Fusion HD cloning kit (TAKARA).

### Neuronal cell culture

To prepare primary neuron cultures, the following procedures (Itoh et al. 2019^66^) were carried out. The brain was dissected from embryonic day 18 (E18) mice of both genders and placed on ice cold dissection medium (1 x HBSS, 1 mM sodium pyruvate, 10 mM HEPES, 30 mM glucose, and 1% penicillin-streptomycin). Hippocampi and cortex were dissected under a binocular microscope and digested with papain (0.5 mg/ml) and DNase (0.01%) in the dissection medium for 25 min at 37 °C. For Dynamin 1 knockout experiments, *Dnm1^+/−^* mice were bred to obtain *Dnm1^+/+^* and *Dnm1^−/−^* animals. Genotyping was immediately performed on clipped tails while brains were kept in dissection medium on ice. Tail DNA was extracted in 50 mM NaCl at 100 °C for 30 min incubation followed by adjusting pH and PCR reaction using custom designed primers (Imoto et al. 2022^33^) and KOD Hot Start DNA Polymerase. For immunocytochemistry, 1.5 x10^5^ neurons were seeded onto 18 mm diameter 1.5H thick coverslips (Neuvitro Corporation) coated with poly-L-lysine (1 mg/mL) in 12-well plates. For Western blots, 6 x10^5^ neurons were seeded on poly-L-lysine coated 6-well plates. Neurons were cultured in Neurobasal media (Gibco) supplemented with 2 mM GlutaMax, 2% B27, 5% horse serum and 1% penicillin-streptomycin at 37 °C in 5% CO_2_. The following day, medium was switched to Neurobasal with 2 mM GlutaMax and 2% B27 (NM0), and neurons were maintained thereafter in this medium. Half of the medium was refreshed every week.

### Lentivirus production

Lentiviral particles expressing Dyn1xA-Flag or Dyn1xB-Flag were produced by co-transfecting HEK293T cells with the expression construct of interest together with two helper plasmids, pHR-CMV8.2 deltaR and pCMV-VSVG, at a molar ratio of 4:3:2. Transfection was performed using polyethyleneimine. Culture supernatants containing viral particles were collected 3 days post-transfection, and viruses were concentrated 20-fold using Amicon Ultra-15 10K centrifugal filter units (Millipore). The concentrated virus was aliquoted, snap-frozen in liquid nitrogen, and stored at -80 °C until use.

### Western blotting

For Western blot confirmation of Dyn1xA antibody, mixed neuron cultures from the hippocampus and cortex were prepared from E18 wild-type and Dnm1 knockout mice and harvested at DIV 14 and DIV 21. For lentiviral expression experiments, mixed neuron cultures from Dnm1 knockout mice were infected at DIV 6 with viruses encoding Dyn1xA-Flag or Dyn1xB-Flag, and cells were harvested at DIV 14. Astrocytes were prepared from the cortex of E 18 wild-type mice and harvested after two weeks. Cells were homogenized in RIPA buffer (Cell Signaling Technology) supplemented with cOmplete Mini Protease Inhibitor (Roche), and centrifuged at 14,000 x g for 10 min at +4 °C. Extracts were separated by SDS–PAGE and transferred onto Immobilon-FL membranes (Millipore Sigma). The membranes were blocked with 3% Bovine Serum Albumin (BSA) in PBS containing 0.05% Tween 20 (PBST) for 1 h, incubated overnight at +4 °C with primary antibodies in 3% BSA in PBST, and then incubated with second antibodies for 1 h at room temperature in 3% BSA in PBST and washed with PBST. The primary antibodies used were: Dynamin1xA (1:2,000), pan-Dynamin1/2/3 (1:2,000), Flag (1:2,000) and β-actin (1:2,000; rabbit). The secondary antibodies were: IRDye 800CW Goat anti-Rabbit IgG (1:4,000) and IRDye 800CW Goat anti-Mouse IgG (1:10,000). Immunocomplexes were detected using an Odyssey infrared imaging system (Li-COR).

For additional Western blot confirmation of Dyn1xA antibody, whole brain lysates from a 1-year-old female wild-type mouse and human not affected with epilepsy were tested (Human Brain Whole Tissue Lysate (Adult Whole Normal), Novus Biologicals). Samples were run at 80 V, 90 min and transferred at 100 V, 80 min. The membrane was blocked with 5% milk in 0.05% PBST for 30 min, incubated overnight at +4 °C with primary antibodies in 5% milk/PBST, and then incubated with second antibodies for 1 h at room temperature in 0.05% PBST before being washed with PBST. The primary antibodies used were Dynamin1xA (1:5,000) and β-actin (1:2,000; mouse). The secondary antibodies were IRDye 800CW Goat anti-Mouse IgG and IRDye 680RD Goat anti-Rabbit IgG (both 1:30,000). Immunocomplexes were detected using an Odyssey infrared imaging system (Li-COR). Precision Plus Protein Dual Color Standard was used as a molecular weight ladder.

### Immunocytochemistry

For immunocytochemistry used to validate the Dyn1xA antibody, DIV14 neurons were fixed with pre-warmed 4% PFA and 4% sucrose in PBS for 20 min, then permeabilized with 0.2% Triton X-100 in PBS for 8 min at room temperature. After blocking with 1% BSA in PBS for 30 min, cells were incubated with primary antibodies in 1% BSA/PBS overnight at 4 °C, followed by secondary antibodies for 1 h at room temperature. Coverslips were mounted with ProLong Diamond Antifade Mountant. Primary antibodies included Dynamin1xA (1:500) and FLAG (1:500). Secondary antibodies were goat anti-rabbit Alexa Fluor 647 (1:500) and goat anti-mouse Alexa Fluor 488 (1:500). Images were acquired using a 60x oil objective lens and the confocal mode tiling function of an Abberior FACILITY line microscope.

## Resource availability

All the materials and protocols will be shared by the lead contact (Shigeki Watanabe) upon request. Raw images in the manuscript will be uploaded to figshare.com upon publication.

## References

1. Moradi Chameh H, Rich S, Wang L, et al. Diversity amongst human cortical pyramidal neurons revealed via their sag currents and frequency preferences. Nat Commun. 2021;12:2497. doi:10.1038/s41467-021-22741-9

2. Barzó P, Szöts I, Tóth M, Csajbók ÉA, Molnár G, Tamás G. Electrophysiology and Morphology of Human Cortical Supragranular Pyramidal Cells in a Wide Age Range. eLife. 2024;13. doi:10.7554/eLife.100390.1

3. Hunt S, Leibner Y, Mertens EJ, et al. Strong and reliable synaptic communication between pyramidal neurons in adult human cerebral cortex. Cereb Cortex. 2022;33(6):2857–2878. doi:10.1093/cercor/bhac246

4. Watson JF, Vargas-Barroso V, Morse-Mora RJ, et al. Human hippocampal CA3 uses specific functional connectivity rules for efficient associative memory. Cell. 2024;0(0). doi:10.1016/j.cell.2024.11.022

5. Nahirney PC, Tremblay ME. Brain Ultrastructure: Putting the Pieces Together. Frontiers in Cell and Developmental Biology. 2021;9. Accessed January 1, 2024. https://www.frontiersin.org/articles/10.3389/fcell.2021.629503

6. St-Pierre MK, Carrier M, González Ibáñez F, et al. Astrocytes display ultrastructural alterations and heterogeneity in the hippocampus of aged APP-PS1 mice and human post-mortem brain samples. Journal of Neuroinflammation. 2023;20(1):73. doi:10.1186/s12974-023-02752-7

7. Oost W, Huitema AJ, Kats K, et al. Pathological ultrastructural alterations of myelinated axons in normal appearing white matter in progressive multiple sclerosis. Acta Neuropathologica Communications. 2023;11(1):100. doi:10.1186/s40478-023-01598-7

8. Shapson-Coe A, Januszewski M, Berger DR, et al. A petavoxel fragment of human cerebral cortex reconstructed at nanoscale resolution. Science. 2024;384(6696):eadk4858. doi:10.1126/science.adk4858

9. Wu JY, Cho SJ, Descant K, et al. Mapping of neuronal and glial primary cilia contactome and connectome in the human cerebral cortex. Neuron. 2024;112(1):41–55.e3. doi:10.1016/j.neuron.2023.09.032

10. Loomba S, Straehle J, Gangadharan V, et al. Connectomic comparison of mouse and human cortex. Science. 2022;377(6602):eabo0924. doi:10.1126/science.abo0924

11. Ultrafast endocytosis at Caenorhabditis elegans neuromuscular junctions | eLife. Accessed November 25, 2024. https://elifesciences.org/articles/00723

12. Watanabe S, Rost BR, Camacho-Pérez M, et al. Ultrafast endocytosis at mouse hippocampal synapses. Nature. 2013;504(7479):242–247. doi:10.1038/nature12809

13. Borges-Merjane C, Kim O, Jonas P. Functional Electron Microscopy, “Flash and Freeze,” of Identified Cortical Synapses in Acute Brain Slices. Neuron. 2020;105(6):992–1006.e6. doi:10.1016/j.neuron.2019.12.022

14. Imig C, López-Murcia FJ, Maus L, et al. Ultrastructural Imaging of Activity-Dependent Synaptic Membrane-Trafficking Events in Cultured Brain Slices. Neuron. 2020;108(5):843–860.e8. doi:10.1016/j.neuron.2020.09.004

15. Andrews JP, Geng J, Voitiuk K, et al. Multimodal evaluation of network activity and optogenetic interventions in human hippocampal slices. Nat Neurosci. 2024;27(12):2487–2499. doi:10.1038/s41593-024-01782-5

16. Schwarz N, Uysal B, Welzer M, et al. Long-term adult human brain slice cultures as a model system to study human CNS circuitry and disease. Slutsky I, Marder E, Mansvelder HD, eds. eLife. 2019;8:e48417. doi:10.7554/eLife.48417

17. Humpel C. Organotypic brain slice cultures: A review. Neuroscience. 2015;305:86–98. doi:10.1016/j.neuroscience.2015.07.086

18. Kusick GF, Chin M, Watanabe S. Zap-and-Freeze Electron Microscopy Captures Synaptic Vesicle Exocytosis with Unprecedented Temporal Precision. Microsc Microanal. 2018;24(S1):1254–1255. doi:10.1017/S143192761800675X

19. Kusick GF, Chin M, Raychaudhuri S, et al. Synaptic vesicles transiently dock to refill release sites. Nat Neurosci. 2020;23(11):1329–1338. doi:10.1038/s41593-020-00716-1

20. Rushton WAH. The effect upon the threshold for nervous excitation of the length of nerve exposed, and the angle between current and nerve. J Physiol. 1927;63(4):357–377.

21. Rudin DO, Eisenman G. THE ACTION POTENTIAL OF SPINAL AXONS IN VITRO. J Gen Physiol. 1954;37(4):505–538.

22. Hoppa MB, Gouzer G, Armbruster M, Ryan TA. Control and Plasticity of the Presynaptic Action Potential Waveform at Small CNS Nerve Terminals. Neuron. 2014;84(4):778–789. doi:10.1016/j.neuron.2014.09.038

23. Atluri PP, Regehr WG. Determinants of the Time Course of Facilitation at the Granule Cell to Purkinje Cell Synapse. J Neurosci. 1996;16(18):5661–5671. doi:10.1523/JNEUROSCI.16-18-05661.1996

24. Lippmann K, Kamintsky L, Kim SY, et al. Epileptiform activity and spreading depolarization in the blood–brain barrier-disrupted peri-infarct hippocampus are associated with impaired GABAergic inhibition and synaptic plasticity. J Cereb Blood Flow Metab. 2017;37(5):1803–1819. doi:10.1177/0271678X16652631

25. Wagner W, Lippmann K, Heisler FF, et al. Myosin VI Drives Clathrin-Mediated AMPA Receptor Endocytosis to Facilitate Cerebellar Long-Term Depression. Cell Reports. 2019;28(1):11–20.e9. doi:10.1016/j.celrep.2019.06.005

26. Lippmann K. A Reduction in the Readily Releasable Vesicle Pool Impairs GABAergic Inhibition in the Hippocampus after Blood–Brain Barrier Dysfunction. Int J Mol Sci. 2024;25(13):6862. doi:10.3390/ijms25136862

27. Bodor AL, Schneider-Mizell CM, Zhang C, et al. The synaptic architecture of layer 5 thick tufted excitatory neurons in mouse visual cortex. Nat Neurosci. 2025;28(8):1704–1715. doi:10.1038/s41593-025-02004-2

28. Mohan H, Verhoog MB, Doreswamy KK, et al. Dendritic and Axonal Architecture of Individual Pyramidal Neurons across Layers of Adult Human Neocortex. Cereb Cortex. 2015;25(12):4839–4853. doi:10.1093/cercor/bhv188

29. Ting JT, Lee BR, Chong P, et al. Preparation of Acute Brain Slices Using an Optimized N-Methyl-D-glucamine Protective Recovery Method. JoVE. 2018;(132):53825. doi:10.3791/53825

30. Watanabe S, Mamer LE, Raychaudhuri S, et al. Synaptojanin and Endophilin Mediate Neck Formation during Ultrafast Endocytosis. Neuron. 2018;98(6):1184-1197.e6. doi:10.1016/j.neuron.2018.06.005

31. Ogunmowo TH, Jing H, Raychaudhuri S, et al. Membrane compression by synaptic vesicle exocytosis triggers ultrafast endocytosis. Nat Commun. 2023;14(1):2888. doi:10.1038/s41467-023-38595-2

32. Imoto Y, Xue J, Luo L, et al. Dynamin 1xA interacts with Endophilin A1 via its spliced long C-terminus for ultrafast endocytosis. The EMBO Journal. 2024;43(16):3327–3357. doi:10.1038/s44318-024-00145-x

33. Imoto Y, Raychaudhuri S, Ma Y, et al. Dynamin is primed at endocytic sites for ultrafast endocytosis. Neuron. 2022;110(17):2815–2835.e13. doi:10.1016/j.neuron.2022.06.010

34. Watanabe S, Trimbuch T, Camacho-Pérez M, et al. Clathrin regenerates synaptic vesicles from endosomes. Nature. 2014;515(7526):228-233. doi:10.1038/nature13846

35. Watanabe S, Liu Q, Davis MW, et al. Ultrafast endocytosis at Caenorhabditis elegans neuromuscular junctions. eLife. 2013;2:e00723. doi:10.7554/eLife.00723

36. Griswold JM, Bonilla-Quintana M, Pepper R, et al. Membrane mechanics dictate axonal pearls-on-a-string morphology and function. Nat Neurosci. Published online December 2, 2024:1–13. doi:10.1038/s41593-024-01813-1

37. Macia E, Ehrlich M, Massol R, Boucrot E, Brunner C, Kirchhausen T. Dynasore, a Cell-Permeable Inhibitor of Dynamin. Developmental Cell. 2006;10(6):839–850. doi:10.1016/j.devcel.2006.04.002

38. Park RJ, Shen H, Liu L, Liu X, Ferguson SM, De Camilli P. Dynamin triple knockout cells reveal off target effects of commonly used dynamin inhibitors. J Cell Sci. 2013;126(22):5305–5312. doi:10.1242/jcs.138578

39. Ceccarelli B, Hurlbut WP, Mauro A. DEPLETION OF VESICLES FROM FROG NEUROMUSCULAR JUNCTIONS BY PROLONGED TETANIC STIMULATION. J Cell Biol. 1972;54(1):30–38.

40. Ge L, Shin W, Arpino G, et al. Sequential compound fusion and kiss-and-run mediate exo- and endocytosis in excitable cells. Sci Adv. 8(24):eabm6049. doi:10.1126/sciadv.abm6049

41. Temporal coincidence between synaptic vesicle fusion and quantal secretion of acetylcholine. J Cell Biol. 1985;101(4):1386–1399.

42. Imoto Y, Xue J, Luo L, et al. Dynamin 1xA interacts with Endophilin A1 via its spliced long C-terminus for ultrafast endocytosis. bioRxiv. Preprint posted online September 24, 2023:2023.09.21.558797. doi:10.1101/2023.09.21.558797

43. Hong Y, Yang Q, Song H, Ming G li. Opportunities and limitations for studying neuropsychiatric disorders using patient-derived induced pluripotent stem cells. Mol Psychiatry. 2023;28(4):1430–1439. doi:10.1038/s41380-023-01990-8

44. Panicker N, Ge P, Dawson VL, Dawson TM. The cell biology of Parkinson’s disease. J Cell Biol. 2021;220(4). doi:10.1083/jcb.202012095

45. Kale MB, Wankhede NL, Bishoyi AK, et al. Emerging biophysical techniques for probing synaptic transmission in neurodegenerative disorders. Neuroscience. 2025;565:63–79. doi:10.1016/j.neuroscience.2024.11.055

46. Herman M, Randall GW, Spiegel JL, Maldonado DJ, Simoes S. Endo-lysosomal dysfunction in neurodegenerative diseases: opinion on current progress and future direction in the use of exosomes as biomarkers. Philos Trans R Soc Lond B Biol Sci. 379(1899):20220387. doi:10.1098/rstb.2022.0387

47. Shin W, Wei L, Arpino G, et al. Preformed Ω-profile closure and kiss-and-run mediate endocytosis and diverse endocytic modes in neuroendocrine chromaffin cells. Neuron. 2021;109(19):3119–3134.e5. doi:10.1016/j.neuron.2021.07.019

48. Calovi S, Soria FN, Tønnesen J. Super-resolution STED microscopy in live brain tissue. Neurobiology of Disease. 2021;156:105420. doi:10.1016/j.nbd.2021.105420

49. Heuser JE, Reese TS. EVIDENCE FOR RECYCLING OF SYNAPTIC VESICLE MEMBRANE DURING TRANSMITTER RELEASE AT THE FROG NEUROMUSCULAR JUNCTION. J Cell Biol. 1973;57(2):315–344.

50. Fesce R, Grohovaz F, Valtorta F, Meldolesi J. Neurotransmitter release: fusion or ‘kiss-and-run’? Trends in Cell Biology. 1994;4(1):1–4. doi:10.1016/0962-8924(94)90025-6

51. Chanaday NL, Cousin MA, Milosevic I, Watanabe S, Morgan JR. The Synaptic Vesicle Cycle Revisited: New Insights into the Modes and Mechanisms. J Neurosci. 2019;39(42):8209–8216. doi:10.1523/JNEUROSCI.1158-19.2019

52. Von Gersdorff H, Mathews G. Dynamics of synaptic vesicle fusion and membrane retrieval in synaptic terminals. Nature. 1994;367(6465):735–739. doi:10.1038/367735a0

53. Soykan T, Kaempf N, Sakaba T, et al. Synaptic Vesicle Endocytosis Occurs on Multiple Timescales and Is Mediated by Formin-Dependent Actin Assembly. Neuron. 2017;93(4):854–866.e4. doi:10.1016/j.neuron.2017.02.011

54. Delvendahl I, Vyleta NP, von Gersdorff H, Hallermann S. Fast, Temperature-Sensitive and Clathrin-Independent Endocytosis at Central Synapses. Neuron. 2016;90(3):492–498. doi:10.1016/j.neuron.2016.03.013

55. Chanaday NL, Kavalali ET. Optical detection of three modes of endocytosis at hippocampal synapses. eLife. 7:e36097. doi:10.7554/eLife.36097

56. Myeong J, Stunault MI, Klyachko VA, Ashrafi G. Metabolic regulation of single synaptic vesicle exo- and endocytosis in hippocampal synapses. Cell Rep. 2024;43(5):114218. doi:10.1016/j.celrep.2024.114218

57. He L, Xue L, Xu J, et al. Compound vesicle fusion increases quantal size and potentiates synaptic transmission. Nature. 2009;459(7243):93–97. doi:10.1038/nature07860

58. Ge L, Shin W, Arpino G, et al. Sequential compound fusion and kiss-and-run mediate exo- and endocytosis in excitable cells. Science Advances. 2022;8(24):eabm6049. doi:10.1126/sciadv.abm6049

59. Galakhova AA, Hunt S, Wilbers R, et al. Evolution of cortical neurons supporting human cognition. Trends in Cognitive Sciences. 2022;26(11):909–922. doi:10.1016/j.tics.2022.08.012

60. Watanabe S, Davis MW, Kusick GF, Iwasa J, Jorgensen EM. SynapsEM: Computer-Assisted Synapse Morphometry. Front Synaptic Neurosci. 2020;12:584549. doi:10.3389/fnsyn.2020.584549

61. Ferguson SM, Brasnjo G, Hayashi M, et al. A selective activity-dependent requirement for dynamin 1 in synaptic vesicle endocytosis. Science. 2007;316(5824):570–574. doi:10.1126/science.1140621

62. Eguchi K, Velicky P, Hollergschwandtner E, et al. Advantages of Acute Brain Slices Prepared at Physiological Temperature in the Characterization of Synaptic Functions. Front Cell Neurosci. 2020;14. doi:10.3389/fncel.2020.00063

63. Cardona A, Saalfeld S, Schindelin J, et al. TrakEM2 Software for Neural Circuit Reconstruction. PLOS ONE. 2012;7(6):e38011. doi:10.1371/journal.pone.0038011

64. Kruzich E, Phadke RA, Brack A, Stroumbakis D, Infante O, Cruz-Martín A. A pipeline for STED super-resolution imaging and Imaris analysis of nanoscale synapse organization in mouse cortical brain slices. STAR Protoc. 2023;4(4):102707. doi:10.1016/j.xpro.2023.102707

65. Sapoznik E, Chang BJ, Huh J, et al. A versatile oblique plane microscope for large-scale and high-resolution imaging of subcellular dynamics. Lakadamyali M, Akhmanova A, Power R, eds. eLife. 2020;9:e57681. doi:10.7554/eLife.57681

66. Itoh K, Murata D, Kato T, et al. Brain-specific Drp1 regulates postsynaptic endocytosis and dendrite formation independently of mitochondrial division. eLife. 8:e44739. doi:10.7554/eLife.44739

