## Supplemental Figures for "Ultrastructural membrane dynamics of mouse and human cortical synapses"

Figure S1. Eddings, et al.

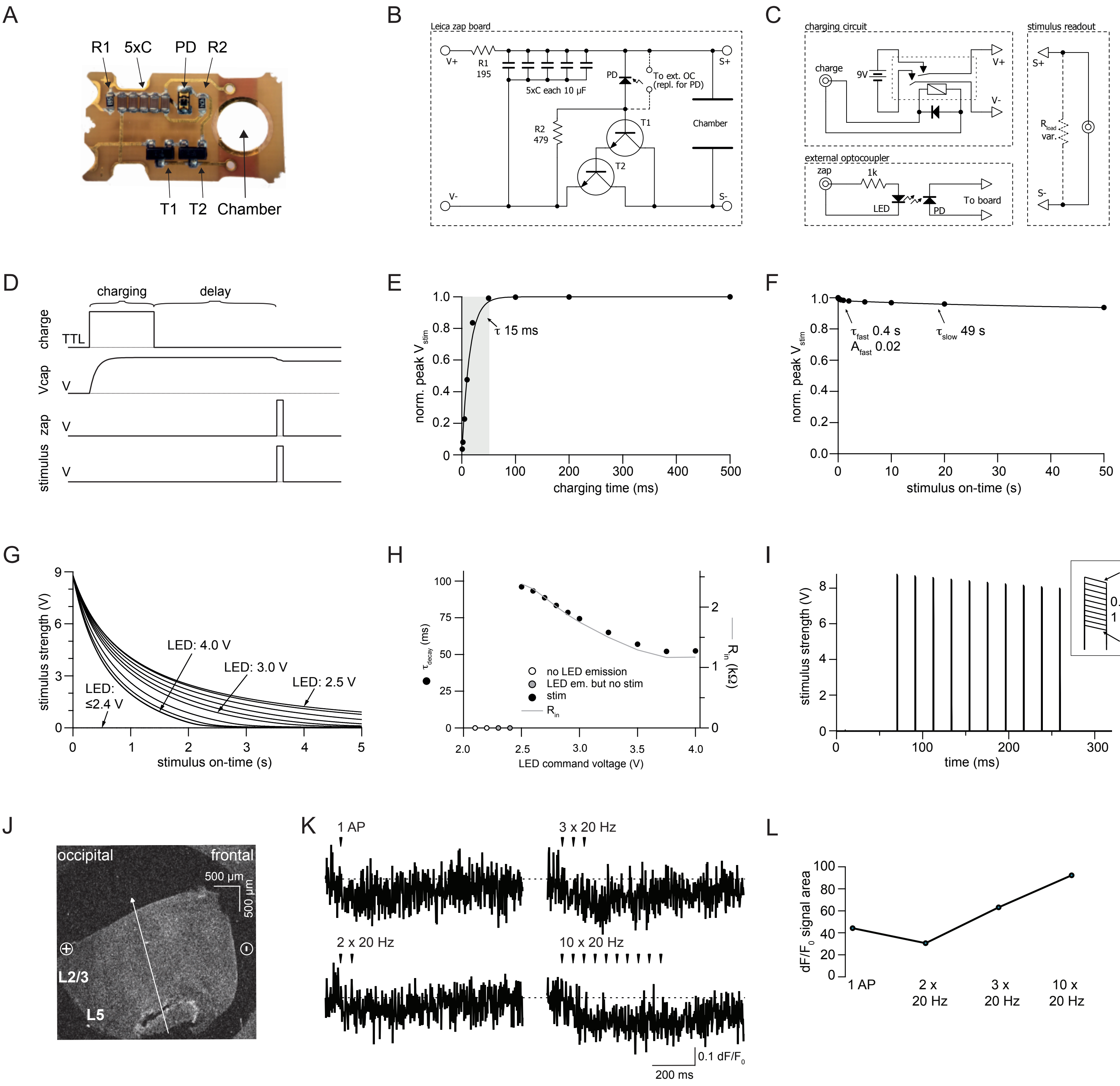

**Figure S1. Electrical circuit of Leica's zap board.**

(A) Leica's zap board. R1: charging resistor, C: capacitor, PD: photodiode, T1 & T2: transistors in Darlington configuration, R2: base-biasing resistor, Chamber: recess for assembly of sapphire disks and Mylar spacer to hold and stimulate the specimen (cf. Figure 1C), V+ and V-: supply voltage (9 V), S+ and S-: stimulus output.

(B) Corresponding equivalent electrical circuit to the zap board. For 2-photon imaging, the PD was removed and replaced by connectors for an external optocoupler (OC, see part C).

(C) Equivalent electrical circuits for charging the capacitors (top left, connected to V+ and V- of the zap board), for the external optocoupler (bottom left, built from a second zap board), and the stimulus read out (right, connected to S+ and S- of the zap board). The charging circuit was controlled by a digital signal (TTL level) to the 'charge' input, the optocoupler via a digital to analog signal to the 'zap' input and electrically driven by a 9 V battery. The 'stimulus readout' was connected to an analog-to-digital converter, with an optional and variable resistor ($R_{load,\ var}$) in parallel. 10 kΩ was used for panel E-I since resistance of a slice ($R_{slice}$) was 12-15 kΩ.

(D) Sequence of signals used to externally operate the zap board. The TTL signal ('charge') activates the relays and allows the 9 V battery to charge the capacitors; 'Vcap' shows the voltage of the capacitors; the 'zap' signal activates the LED, which activates the PD, Darlington transistors and stimulus.

(E) Normalized peak stimulus amplitude (1 ms) versus charging time for $R_{load}$ of 10 kΩ (other R values gave similar results). The gray area denotes the standard charging time used in this figure and Figure 1. The fit was forced to go from 0 to 1.

(F) Normalized peak stimulus amplitude (1 ms stimulus duration) versus delay time for $R_{load}$ of 10 kΩ data, whereas other $R_{load}$ values resulted in similar values. Double-exponential fit was forced to go from 1 to 0. The fast decay time $\tau_{fast}$ is likely due to charging of the analog-to-digital (AD) input (220 pF, HEKA instruments) via the leak current of the PD and Darlington transistors.

(G) Voltage output curves for continuous zap commands. The light intensity shone on the PD was varied from 2.4-4 V supply voltage for the LED (in 100 mV and 250 mV steps up to 3 and 4 V, respectively). Voltage outputs for 3.75 and 4 V overlap, because the LED emission saturated. Failures at 2-2.4 V were omitted for clarity. Data shown for 10 kΩ $R_{load}$. The uneven curve patterns likely result from the nonlinear behavior of the circuit consisting of LED, PD and Darlington transistors.

(H) For each strength of LED illumination ('LED command voltage'), the decay time of a single exponential fit to the data shown in G is plotted (the fit was restricted from the peak to the time at which 2 V was reached, to exclude the nonlinearities expected to arise when voltage across the capacitors dropped below the working range of the Darlington transistors). Failures are plotted as open/grey symbols. The grey line shows the light-dependent resistance of the PD ($R_{PD}$), calculated from $\tau = R*C$, with C = 50 µF and R represented by the resistances of the AD input (1 MΩ, HEKA instruments) in parallel with R2 (479 Ω) plus $R_{PD}$ (in series).

(I) Plot of stimulus strength over a train of 10 stimuli given at 50 Hz ($R_{load}$ = 10kΩ). Inset: zoom-in on the peaks of superimposed stimuli.

(J) A 100 µm sagittal slice of the prefrontal cortex (frontal and occipital orientation indicated) assembled between two sapphire discs for 2-photon imaging on the zap board. Layer 2/3 (L2/3) and layer 5 (L5) are indicated along with an arrow depicting the orientation of the line scan.

(K) Representative normalized Fura-2 AM signals following a single action potential (upper left), two action potentials (lower left), three action potentials (upper right), or ten consecutive stimuli (lower right), each at 20 Hz, from the cortical slice shown in (J). Dotted line indicates baseline at y=1.

(L) The calculated dF/F0 signal area is plotted for the different stimulation paradigms of the cortex example shown in (J, K). The area was calculated between y=1 (dotted line) and the signal trace, starting after the first stimulus at 70 ms and ending at 900 ms.

Figure S2. Eddings, et al.

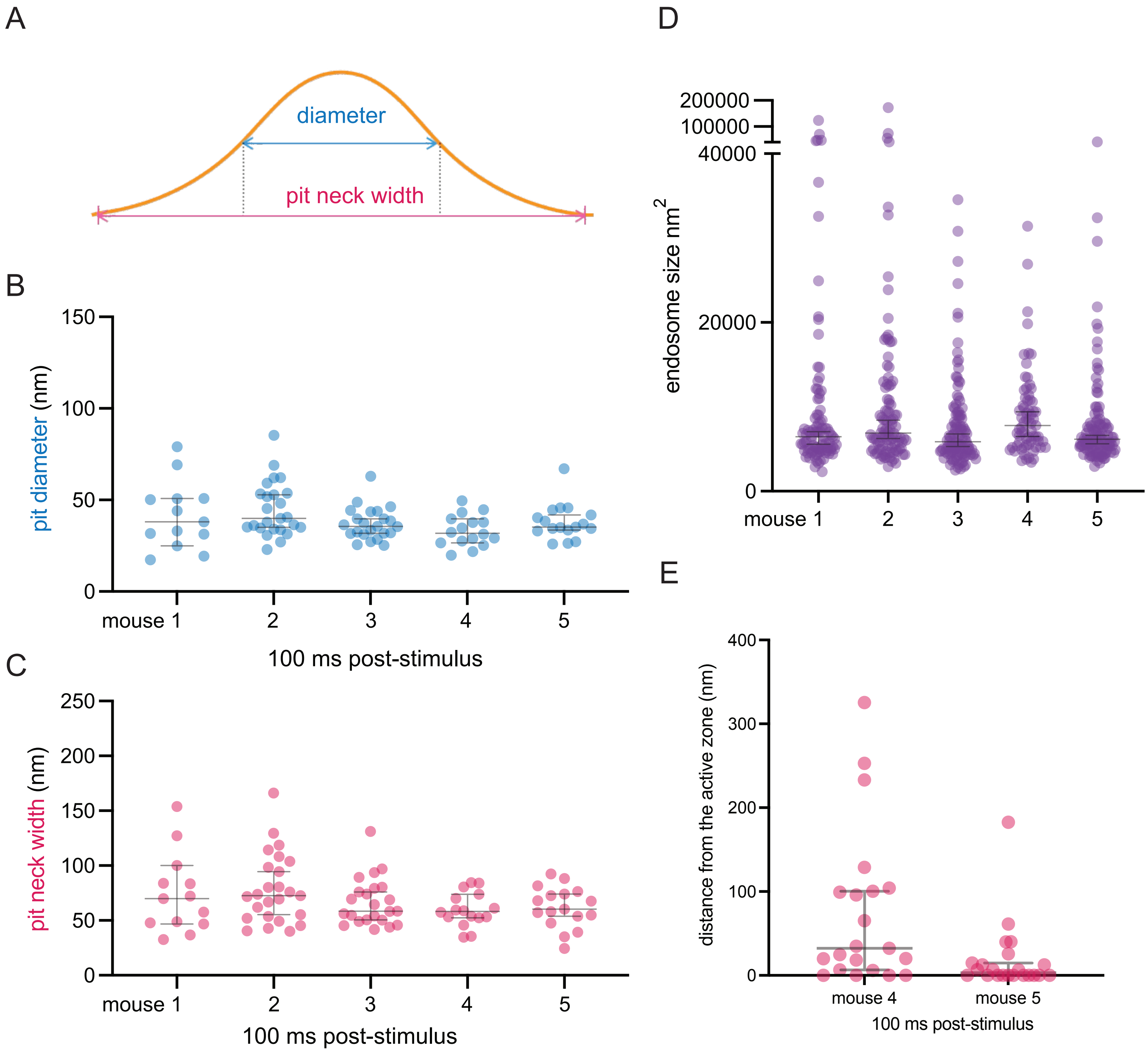

**Figure S2.** Size of uncoated pits and endosomes in acute mouse cortical slices.
(A) Cartoon illustrating the areas that have been measured as the pit diameter (blue) and pit neck width (pink). The diameter is measured at the full-width half maximum.
(B) Plot showing the diameter of uncoated pits 100 ms post-stimulus in acute mouse cortical slices from n=3 mice. Data are presented as median ± 95% confidence interval. Each dot represents a pit.
(C) Plot showing the neck width of uncoated pits 100 ms post-stimulus in acute mouse cortical slices from n=3 mice. Data are presented as median ± 95% confidence interval. Each dot represents a pit.
(D) Plot showing the sizes of endosomes across all tested timepoints in acute mouse cortical slices from n=3 mice. Data are presented as median ± 95% confidence interval. Each dot represents an endosome.
(E) Plot showing the distance distribution of putative uncoated endocytic pits from the edge of an active zone 100 ms post-stimulus in acute slices from n=2 mice (mouse 4 and mouse 5). Data are presented as median ± 95% confidence interval. Each dot represents a pit.

Figure S3. Eddings, et al.

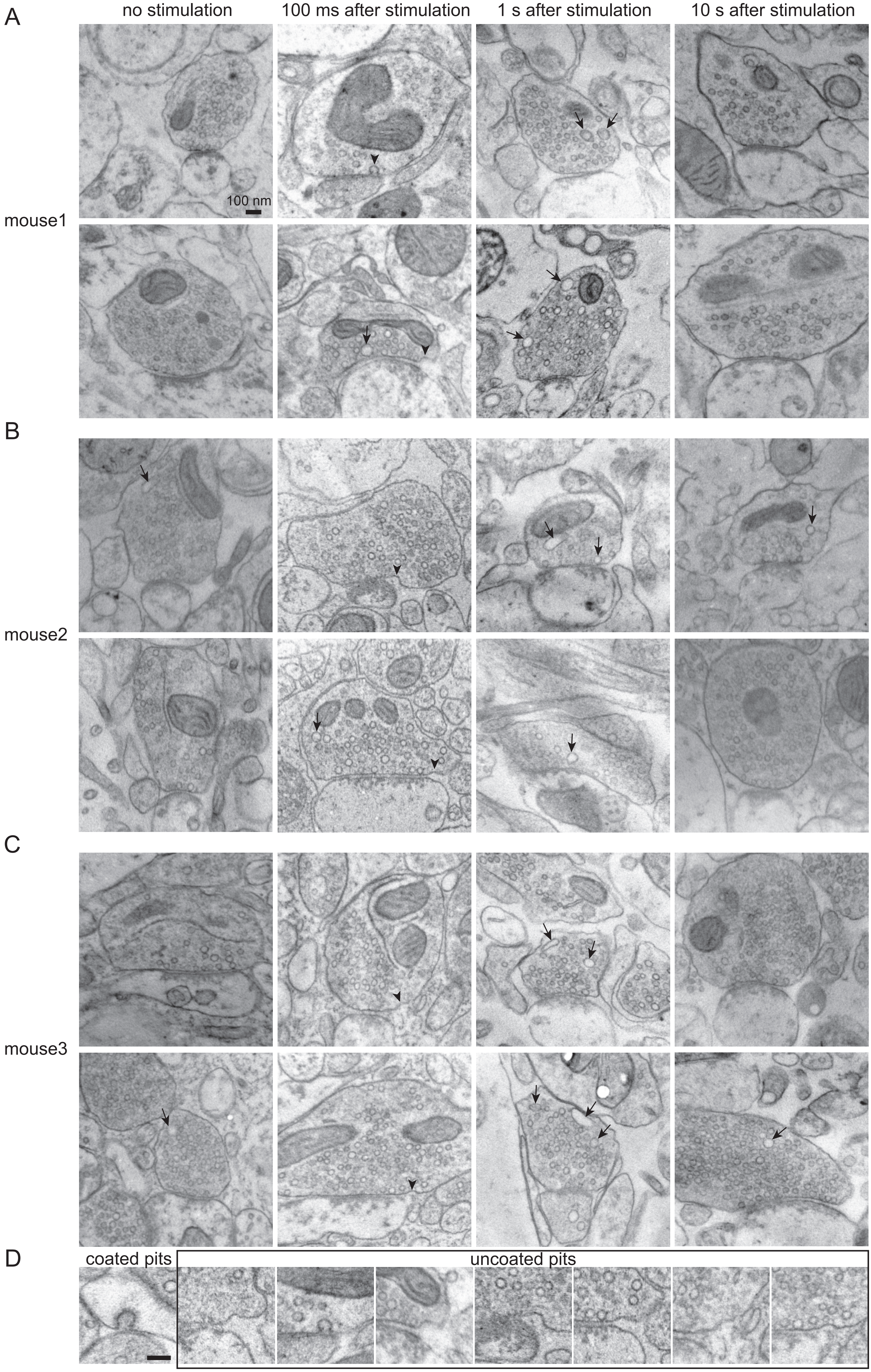

**Figure S3.** Additional EM images for Figure 2.
(A-C) Electron micrographs of acute mouse brain slices that have undergone zap-and-freeze at the indicated time points.
(D) An example clathrin-coated pit is shown compared to uncoated pits from this dataset. Scale bar: 100 nm.

Figure S4. Eddings, et al.

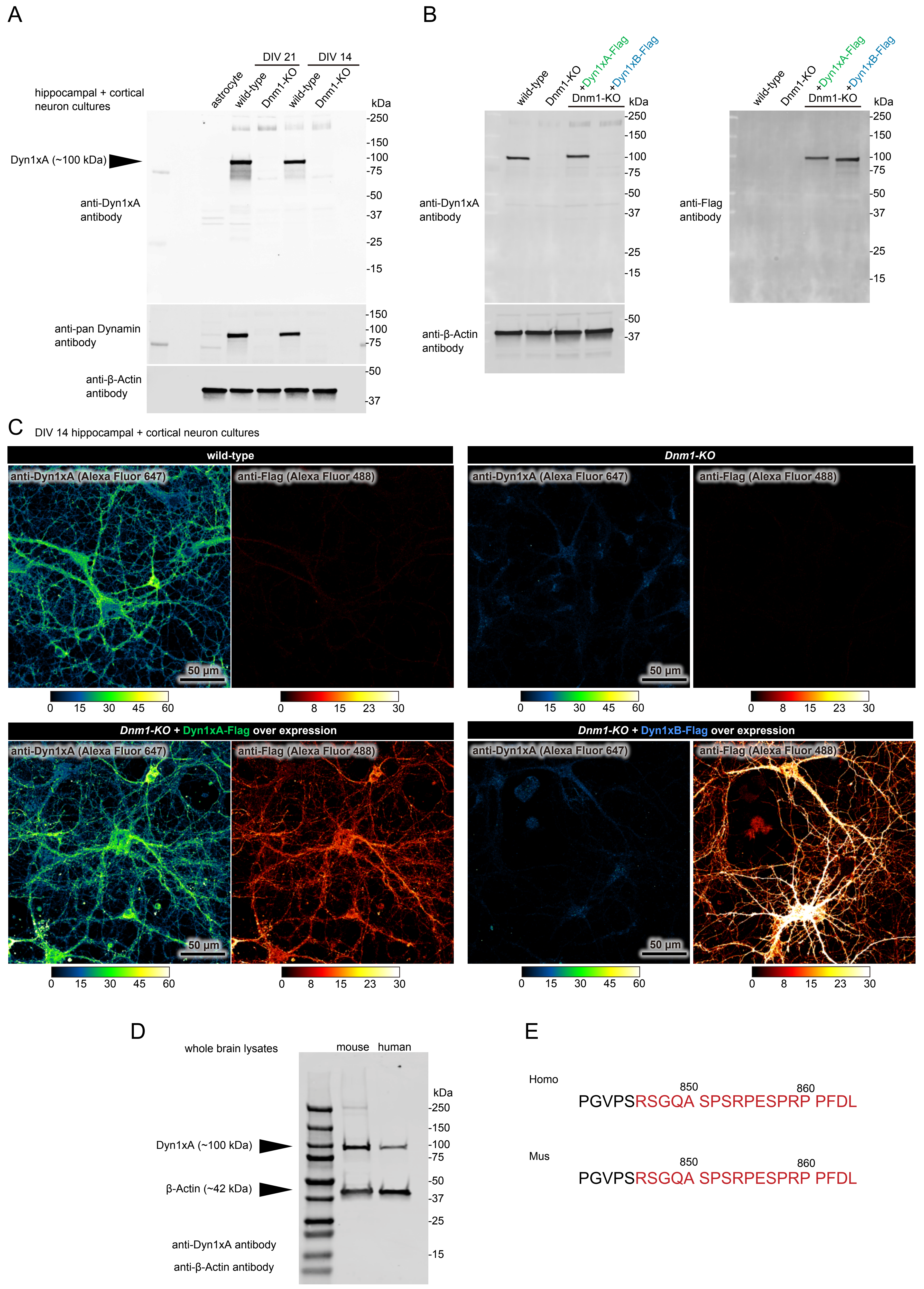

**Figure S4.** Validation of Dynamin1xA antibody.
(A) Western blot showing protein detection using our lab-made anti-Dyn1xA antibody and a commercially available anti-pan Dynamin1/2/3 antibody. Equal amounts (20 µg) of protein from astrocytes and neuron cultures from wild-type and Dynamin 1 (*Dnm1*) knockout mice at DIV 14 and DIV 21 were analyzed. Anti-beta-Actin antibody was used for protein loading control.
(B) Western blot showing specificity of anti-Dyn1xA antibody. Equal amounts (20 µg) of protein from DIV 14 neuron cultures from wild-type, *Dnm1* knockout, and *Dnm1* knockout + lentiviral overexpression of Dyn1xA-Flag or Dyn1xB-Flag were analyzed. Anti-beta-Actin antibody was used for protein loading control. Anti-Flag antibody was used to confirm the expression of exogenous Dyn1xA-Flag and Dyn1xB-Flag.
(C) Immunofluorescence images showing specific detection of Dyn1xA in primary neurons using the lab-made anti-Dyn1xA antibody and anti-Flag antibody as a control. DIV 14 cultured neurons from wild-type, *Dnm1* knockout, and *Dnm1* knockout + lentiviral overexpression of Dyn1xA-Flag or Dyn1xB-Flag were analyzed. Heat scale indicates fluorescence intensity for each imaging channel: darker colors indicate no or low signal, brighter colors indicate higher signal.
(D) Western blot showing anti-Dyn1xA antibody reactivity with whole brain lysates from wild-type mouse and human (note: samples were not loaded with a specific protein concentration). Mouse lysate is from a 1-year-old female wild-type mouse. Human lysate is ‘Human Brain Whole Tissue Lysate (Adult Whole Normal)’ from Novus Biologicals, and not from a tested epilepsy patient used in this study.
(E) Sequence of the Dyn1xA C-terminus in mice and humans. Sequence used to generate the antibody epitope is highlighted in red.

Figure S5. Eddings, et al.

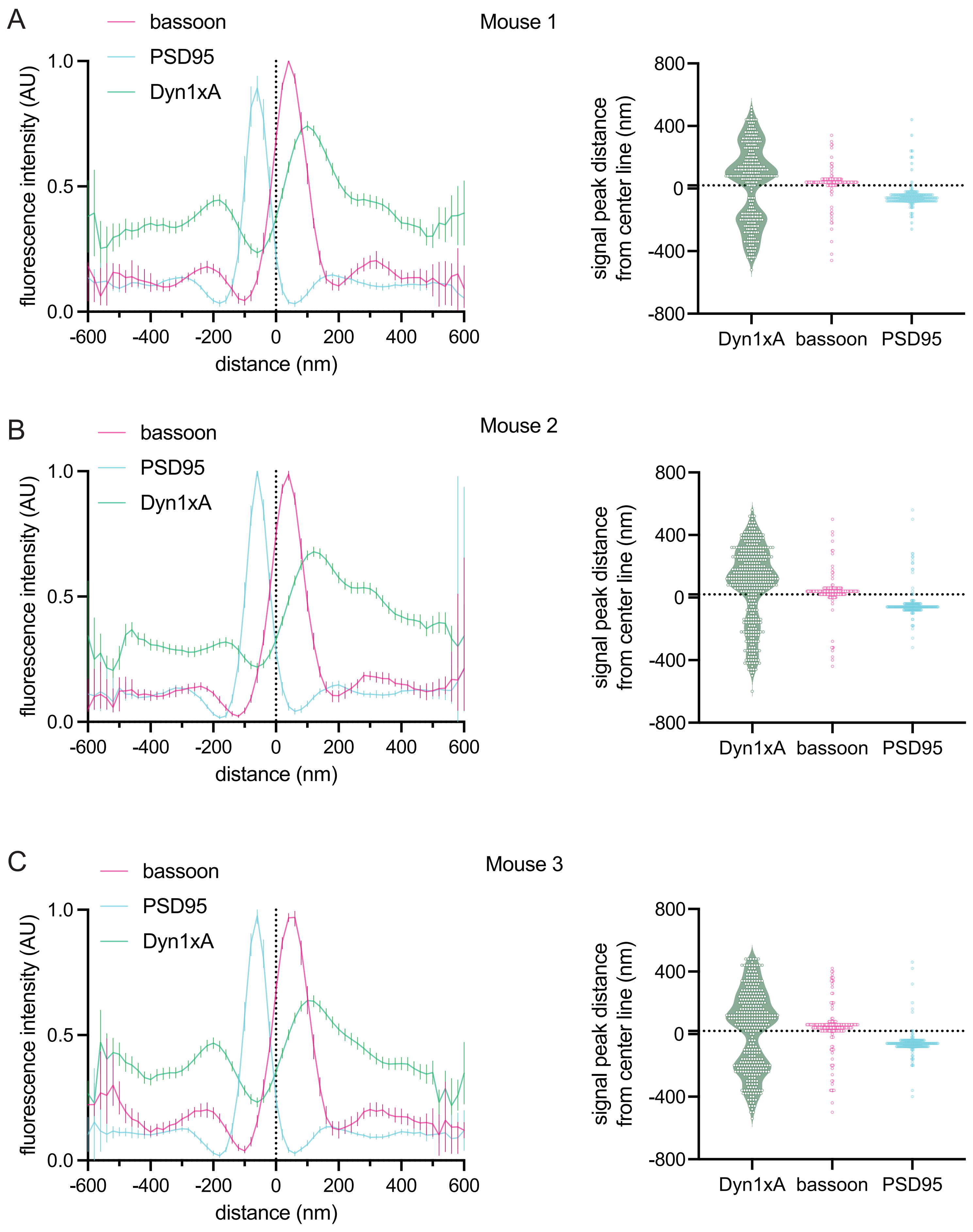

**Figure S5.** Individual traces for Figure 4.
Line scan profiles and individual peak distances obtained from side view synapses for mouse 1 (A), 2 (B), and 3 (C) respectively; see Data Table S1 for specific n values. Dotted line indicates center-line at x=0.

Figure S6. Eddings, et al.

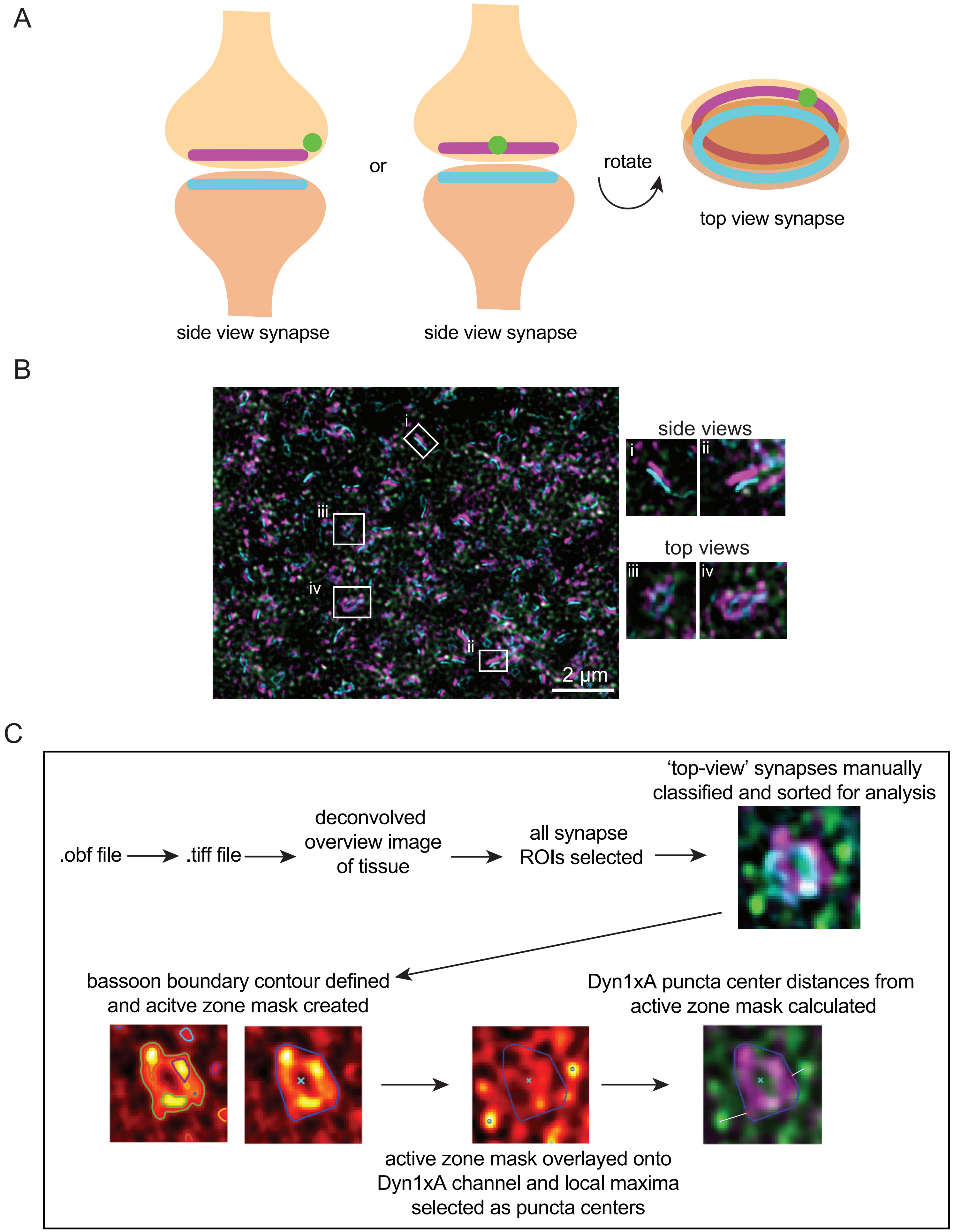

**Figure S6.** Additional mouse STED images for Figure 4.
(A) Cartoon illustrating the orientation of what's been classified as a 'side-view' versus 'top-view' synapse. Magenta: Bassoon presynapse, cyan: PSD95 postsynapse, green: Dyn1xA puncta. Dyn1xA puncta can appear in the middle of Bassoon signals from side-views, but *en face* top-views show Dyn1xA puncta is at the edge of a Bassoon ring.
(B) Example Dyn1xA puncta in side view excitatory synapse images.
(C) Visual workflow showing how STED images are handled in the top view analysis pipeline described in detail in Methods. Starting from .obf raw files obtained from the STED microscope onto how puncta distances are calculated using MATLAB scripts.

Figure S7. Eddings, et al.

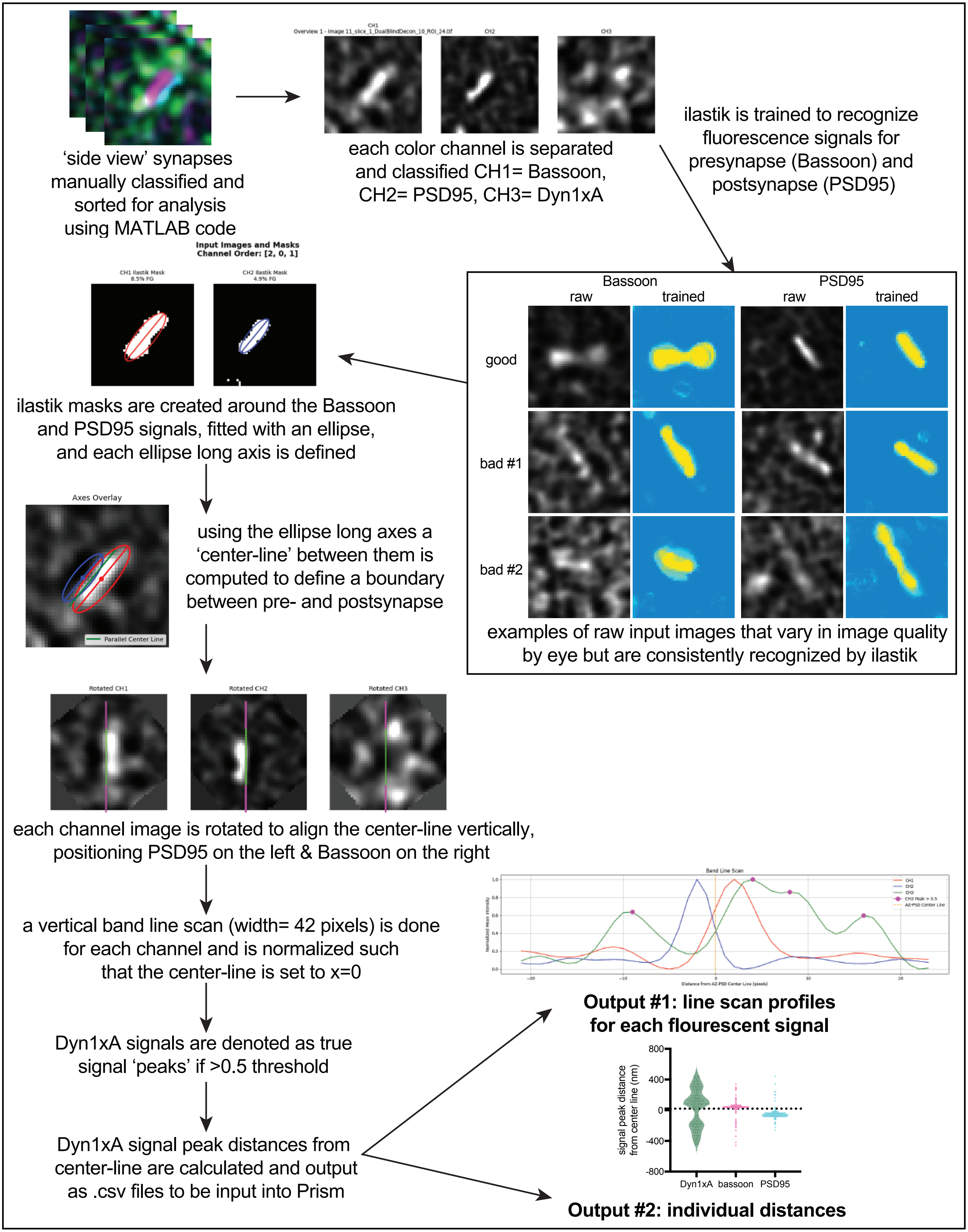

**Figure S7.** STED side view analysis pipeline.
Visual workflow showing how STED side view images are handled in the pipeline detailed in Methods.

Figure S8. Eddings, et al.

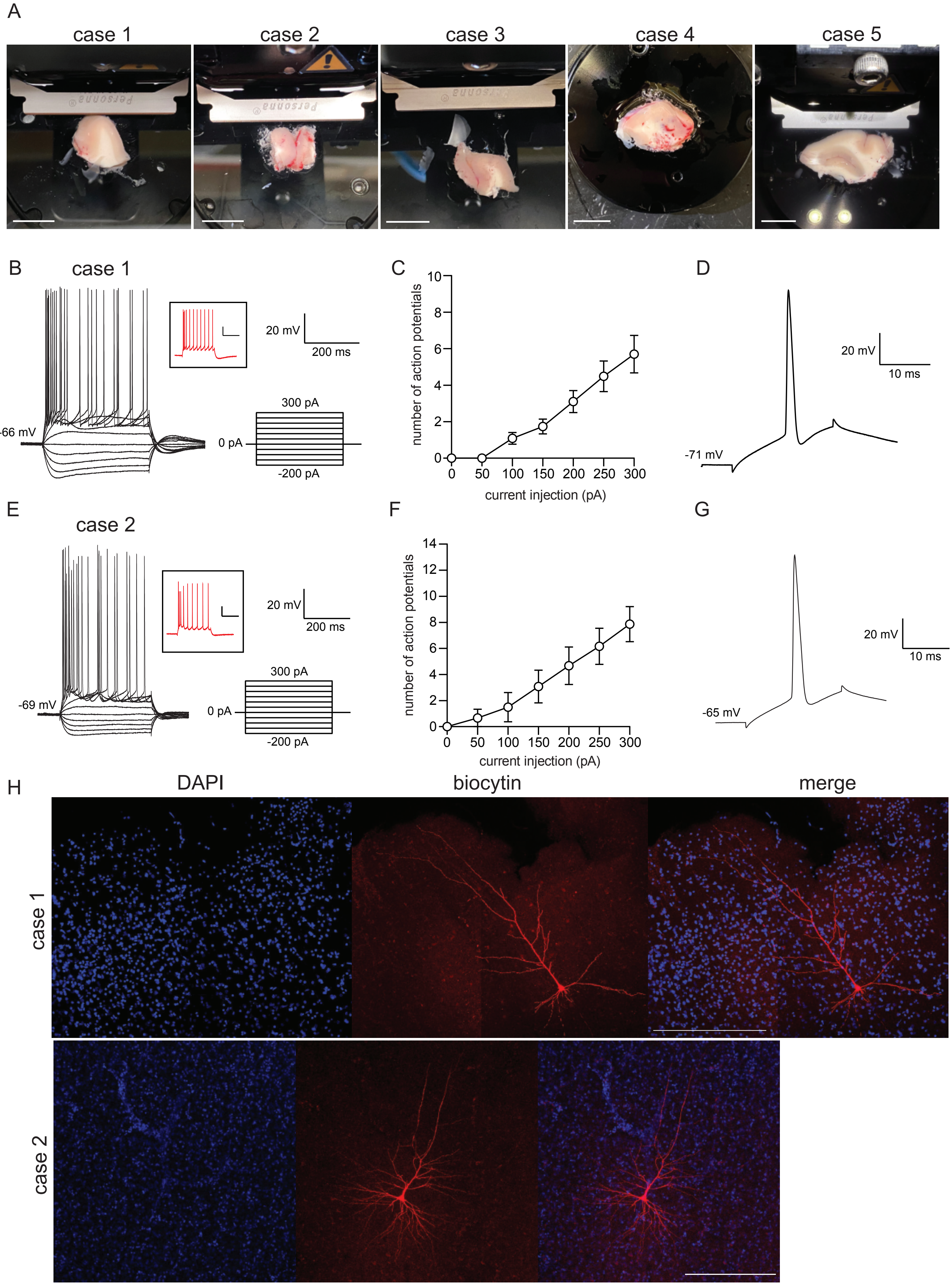

**Figure S8.** Electrophysiological validation of human cortical slice viability after NMDG recovery.
(A) Photos of samples from each case on a vibratome. Scale bars: 1 cm. Note: photo was not obtained for final case 6 sample while it was on the vibratome.
(B) Representative traces from case 1 cortex pyramidal neuron in response to a series of 400 ms current stepping from -200 to +300 pA with increments of 50 pA. Inset: representative trace in response to +300 pA injection.
(C) The average number of action potentials generated in response to depolarizing current pulses. n = 10 cells from 4 slices; case 1.
(D) Typical spike of case 1 human cortical pyramidal neuron obtained at the normal resting membrane potential (RMP).
(E) Representative traces from case 2 cortex pyramidal neuron in response to a series of 400 ms current stepping from -200 to +300 pA with increments of 50 pA. Inset: representative trace in response to +300 pA injection.
(F) The average number of action potentials generated in response to depolarizing current pulses. n = 6 cells from 2 slices; case 2.
(G) Typical spike of case 2 human cortical pyramidal neuron obtained at the normal RMP.
(H) Confocal images of pyramidal neurons filled by biocytin post-recording from case 1 and case 2. Scale bars: 300 μm.

Figure S9. Eddings, et al.

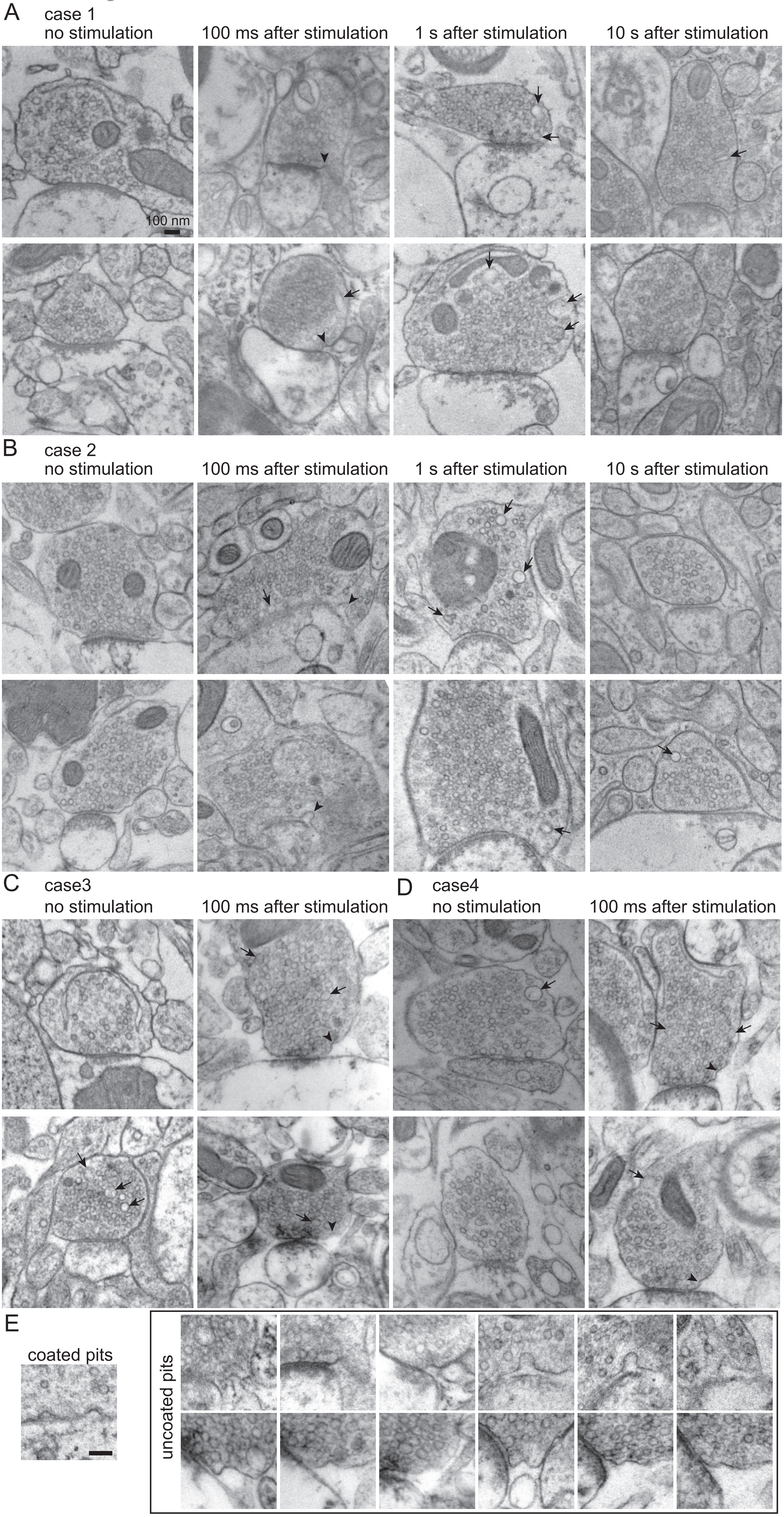

**Figure S9.** Additional EM images for Figure 5.
(A-D) Electron micrographs of acute human brain slices that have undergone zap-and-freeze at the indicated time points.
(E) Example clathrin-coated pits are shown compared to uncoated pits from this dataset. Scale bar: 100 nm.

Figure S10. Eddings, et al.

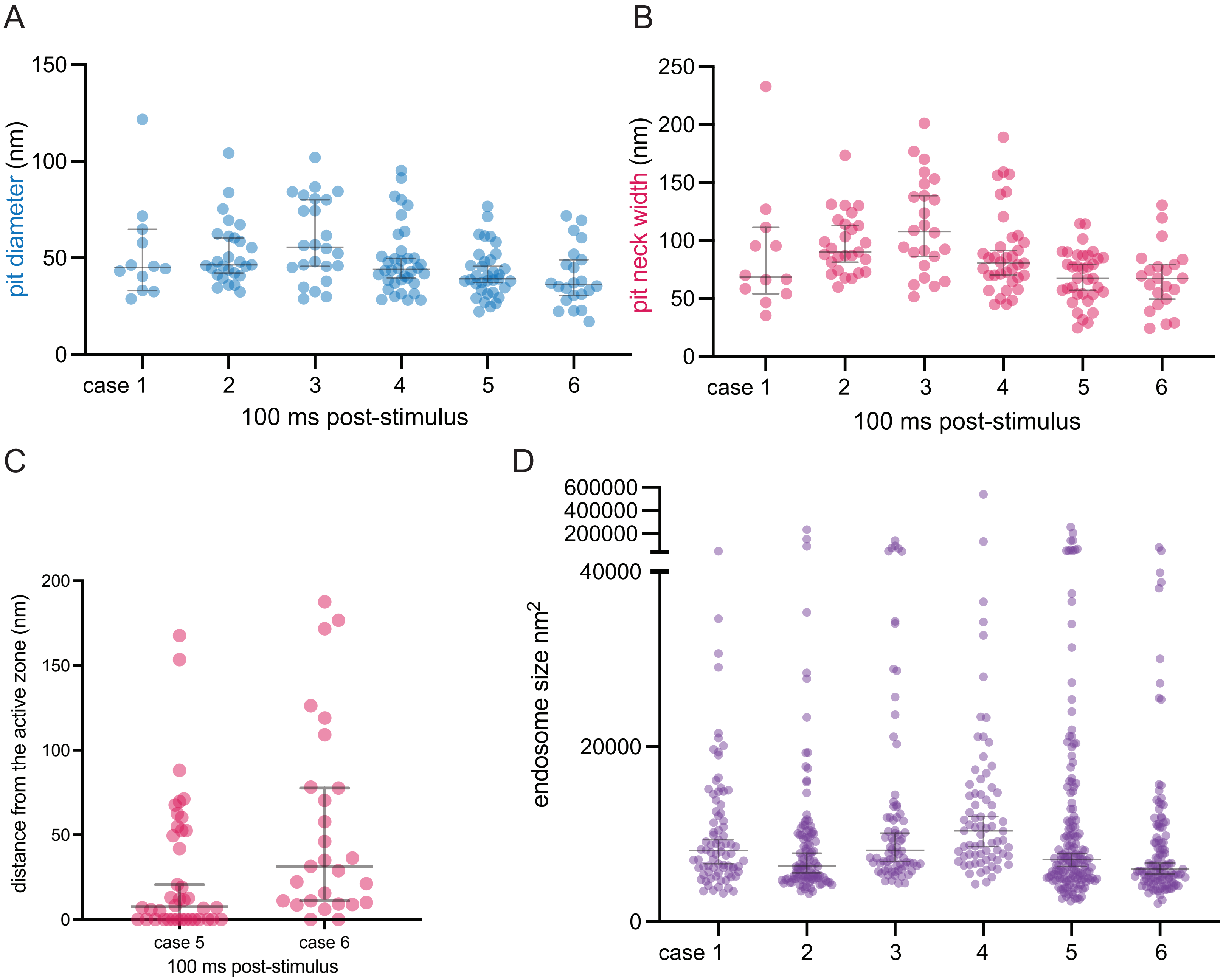

**Figure S10.** Size of uncoated pits and endosomes in acute human cortical slices.
(A) Plot showing the diameter of uncoated pits 100 ms post-stimulus in acute human cortical slices from n=6 humans (case 1-6). Data are presented as median ±95% confidence interval. Each dot represents a pit.
(B) Plot showing the neck width of uncoated pits 100 ms post-stimulus in acute human cortical slices from n=6 humans (case 1-6). Data are presented as median ±95% confidence interval. Each dot represents a pit.
(C) Plot showing the distance distribution of putative uncoated endocytic pits from the edge of an active zone 100 ms post-stimulus in acute neocortical slices from n=2 humans (case 5 and case 6). Data are presented as median ±95% confidence interval. Each dot represents a pit.
(D) Plot showing the sizes of endosomes across all tested timepoints in acute human cortical slices from n=6 humans (case 1-6). Data are presented as median ±95% confidence interval. Each dot represents an endosome.

Figure S11. Eddings, et al.

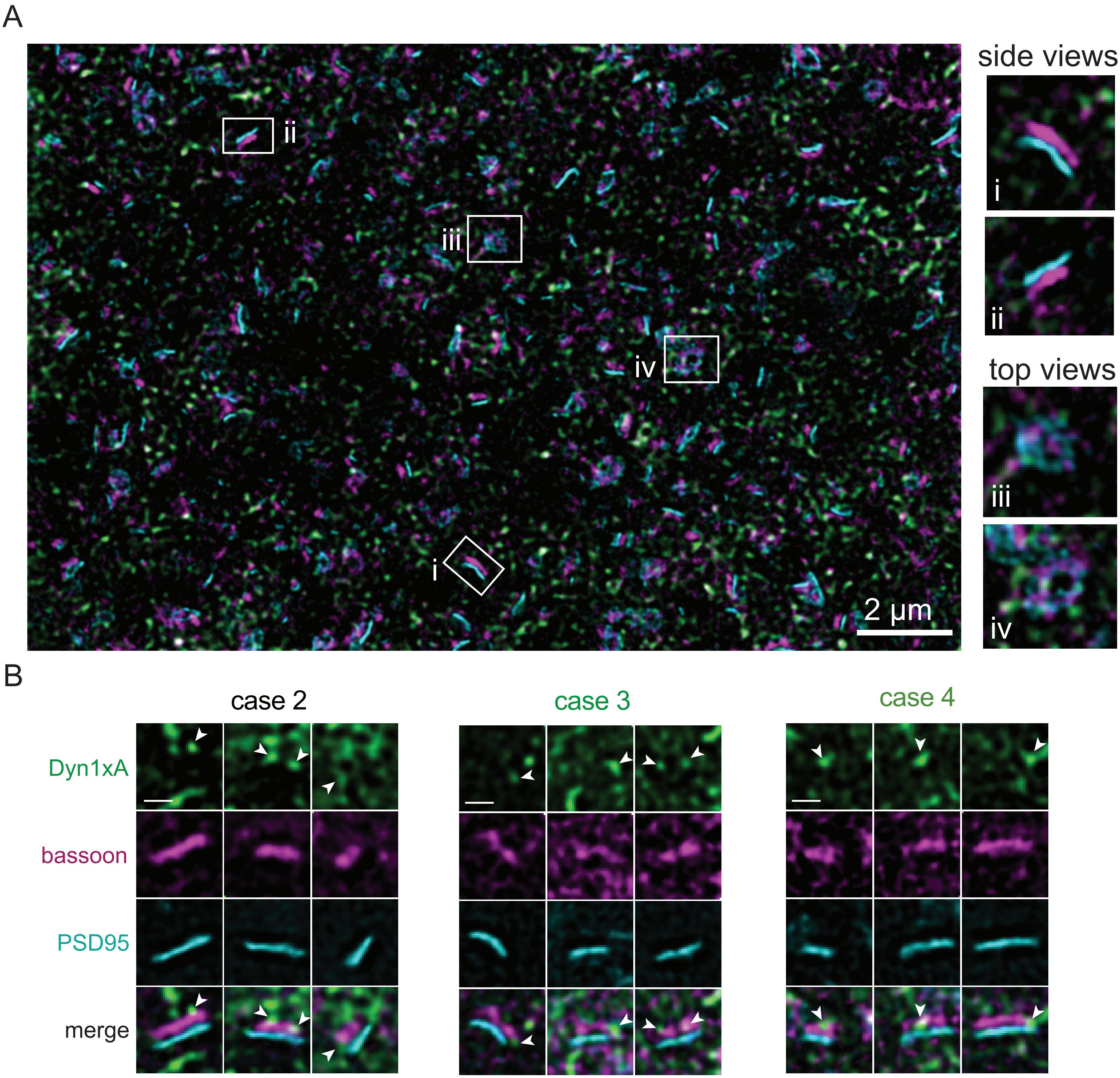

**Figure S11.** Additional human STED images for Figure 7.
(A) Overview 2D, three-color STED image of an acute human neocortical slice (from case 2). Example side view (i and ii) and top view (iii and iv) synapses are highlighted as panels. Scale bar: 300 nm unless noted.
(B) Example Dyn1xA puncta in side view excitatory synapse images. Note: some Dyn1xA puncta appear to be in the middle of Bassoon signals when observed from side views, and this is why top view images were analyzed.

Figure S12. Eddings, et al.

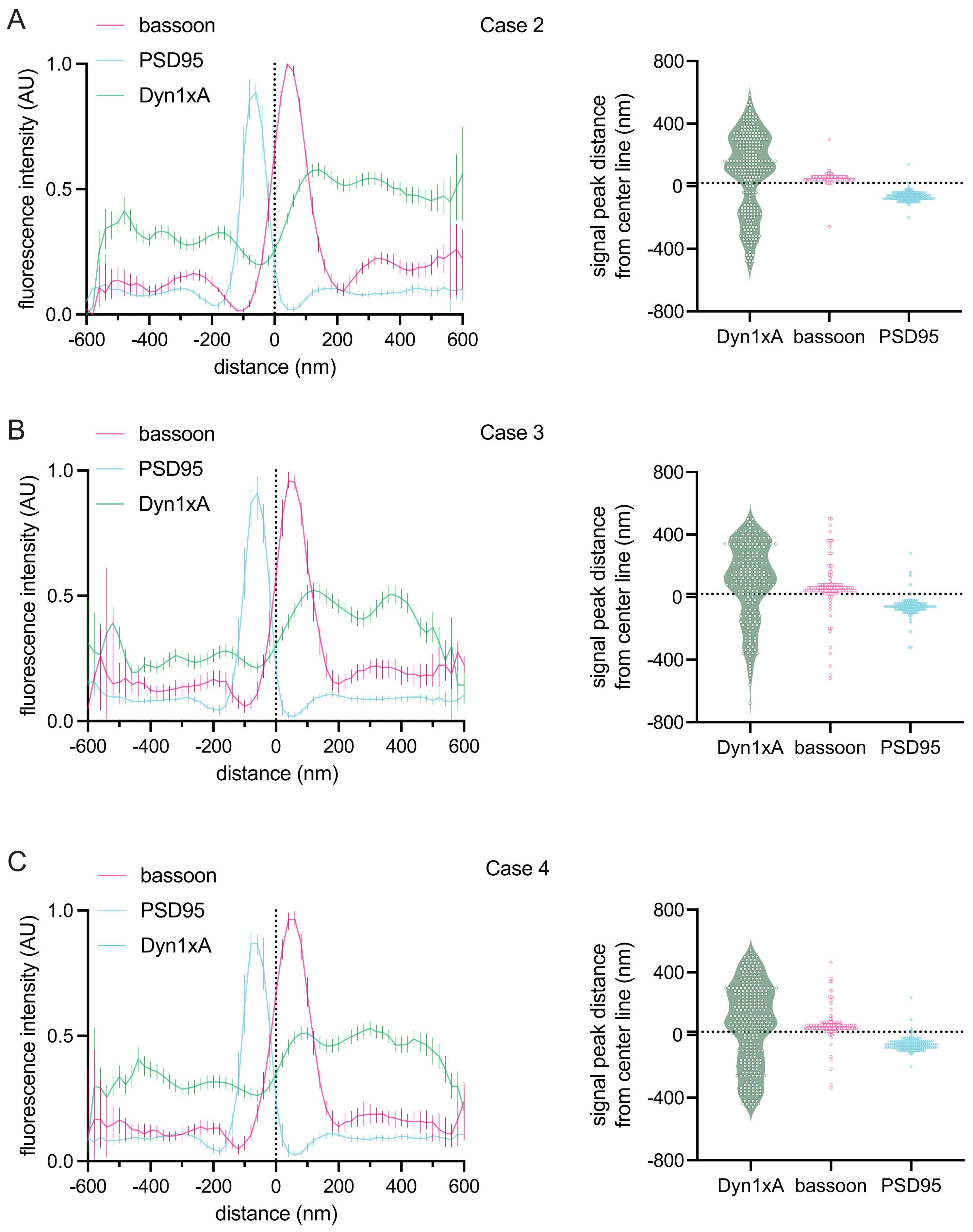

**Figure S12.** Individual traces for Figure 7.
Line scan profiles and individual peak distances obtained from side view synapses for case 2 (A), case 3 (B), and case 4 (C) respectively; see Data Table S1 for specific n values. Dotted line indicates center-line at x=0.
